# *Toxoplasma gondii* excretion of glycolytic products is associated with acidification of the parasitophorous vacuole during parasite egress

**DOI:** 10.1101/2021.11.25.469974

**Authors:** My-Hang Huynh, Vern B. Carruthers

## Abstract

The *Toxoplasma gondii* lytic cycle is a repetition of host cell invasion, replication, egress, and re-invasion into the next host cell. While the molecular players involved in egress have been studied in greater detail in recent years, the signals and pathways for triggering egress from the host cell have not been fully elucidated. A perforin-like protein, PLP1, has been shown to be necessary for permeabilizing the parasitophorous vacuole (PV) membrane or exit from the host cell. *In vitro* studies indicated that PLP1 is most active in acidic conditions, and indirect evidence using superecliptic pHluorin indicated that the PV pH drops prior to parasite egress. Using ratiometric pHluorin, a GFP variant that responds to changes in pH with changes in its bimodal excitation spectrum peaks, allowed us to directly measure the pH in the PV prior to and during egress by live-imaging microscopy. A statistically significant change was observed in PV pH during egress in both wild-type RH and Δ*plp1* vacuoles compared to DMSO-treated vacuoles. Interestingly, if parasites are chemically paralyzed, a pH drop is still observed in RH but not in Δ*plp1* tachyzoites. This indicates that the pH drop is dependent on the presence of PLP1 or motility. Efforts to determine transporters, exchangers, or pumps that could contribute to the drop in PV pH identified two formate-nitrite transporters (FNTs). Auxin-induced conditional knockdown and knockouts of FNT1 and FNT2 reduced the levels of lactate and pyruvate released by the parasites and lead an abatement of vacuolar acidification. While additional transporters and molecules are undoubtedly involved, we provide evidence of a definitive reduction in vacuolar pH associated with induced and natural egress and characterize two transporters that contribute to the acidification.

**Author Summary:** *Toxoplasma gondii* is a single celled intracellular parasite that infects many different animals, and it is thought to infect up to one third of the human population. This parasite must rupture out of its replicative compartment and the host cell to spread from one cell to another. Previous studies indicated that a decrease in pH occurs within the replicative compartment near the time of parasite exit from host cells, an event termed egress. However, it remained unknown whether the decrease in pH is directly tied to egress and, if so, what is responsible for the decrease in pH. Here we used a fluorescent reporter protein to directly measure pH within the replicative compartment during parasite egress. We found that pH decreases immediately prior to parasite egress and that this decrease is linked to parasite disruption of membranes. We also identified a family of transporters that release acidic products from parasite use of glucose for energy as contributing to the decrease in pH during egress. Our findings provide new insight that connects parasite glucose metabolism to acidification of its replicative compartment during egress from infected cells.

## Introduction

As an obligate intracellular pathogen, *Toxoplasma gondii* critically relies on efficiently completing each step of its lytic cycle for successful propagation and survival. In recent years, a greater focus has been applied to the egress step, wherein the parasites exit the host cell as a necessary prelude to infecting a neighboring cell. This increased attention has elucidated a well-orchestrated and complex cascade of events involving several molecular players, including activation of protein kinases by calcium (Ca^2+^) or cyclic guanine monophosphate, release of proteins from apical microneme organelles, and activation of the glideosome motility machinery (1). As with most other parasites belonging to the phylum Apicomplexa, *T. gondii* parasites replicate inside a membrane bound parasitophorous vacuole (PV), from which it must escape before leaving the host cell to initiate another round of the lytic cycle.

Most of the studies involving egress have been performed using chemical inducers such as Ca^2+^ ionophores that increase cytosolic Ca^2+^ to elicit egress. A proposed general model for the activation of egress is as follows (2): addition of inducer increases host and parasite Ca^2+^ levels, which leads to the secretion of the micronemes, including the perforin-like protein 1 (PLP1) and perforation of the PVM. Permeabilization of the PVM causes an influx of Ca^2+^ from the host cell and medium, leading to a second peak of Ca^2+^ increase in the parasites, activation of the motility machinery, exit from the host cell. During spontaneous or natural egress, diacylglycerol kinase 2 (DGK2) was shown to be a plausible candidate for an intrinsic signal (3). DGK2 is secreted into the PV and generates phosphatidic acid as a signaling molecule. DGK2-defective parasites were selectively defective in natural egress but were able to respond to chemical inducers. Conversely, egress is negatively regulated by the cAMP-dependent protein kinase A catalytic subunit 1 (PKAc1) through suppression of cyclic GMP (cGMP) cytosolic Ca^2+^ signaling, as demonstrated by premature egress of PKAc1-deficient parasites (4, 5). These studies also showed that acidification was necessary for the early egress of PKAc1-deficient tachyzoites as determined through treatment with a P-type ATPase inhibitor (dicyclohexylcarbodiimide, DCCD) ((5) or neutralization with NH_4_Cl (4).

Early work showed that tachyzoite motility is pH-dependent, and that alkaline conditions inhibit motility whereas acidic buffers induce motility (6). It was later shown that this pH-dependent motility is likely due to the effect of pH on microneme secretion, wherein low pH activates microneme secretion, and also leads to parasite egress (7). Concomitantly, pH neutralization or treatment with DCCD was found to suppress tachyzoite egress. This study also noted that a drop in PV pH was associated with late-stage infection ∼30 h post-inoculation, a decrease that was partially reversed upon treatment with the weak base NH_4_Cl. The low pH associated with egress was correlated with an increase in PLP1 activity. More specifically, in erythrocyte hemolysis assays, recombinant PLP1 lytic activity was shown to be most active between pH 5.4 and 6.4, a pH range for which PLP1 binding to erythrocyte ghost membranes was also enhanced (7). *T. gondii* PLP1 is not unique in this respect, as several other pathogenic cytolytic proteins have been found to be more active at low pH, including *Listeria monocytogenes* listeriolysin O (8, 9), *Leishmania amazonensis* a-leishporin (10), *Trypanosoma cruzi* TC-TOX (11), and *Bacillus thuringiensis* Cyt1A (12). Also, acidification of transient PVs containing *Plasmodium yoelii* sporozoites was suggested to promote parasite escape in a PyPLP1-dependent manner (13). For TgPLP1, increased activity at low pH is consistent with the observed lower pH of the PV beginning at ∼30 h post-infection when parasites are beginning to egress (7). However, the study was carried out using superecliptic pHluorin, which measures relative pH (14) rather than absolute pH. Also, the measurements were made on a population of infected cells over a relatively long period of time. Thus, this earlier work failed to capture changes in pH occurring during egress from individual cells.

Herein, we utilize a ratiometric pHluorin (RatpH) (14) to directly quantify PV pH in individual infected cells. Expression of RatpH in the PV consistently revealed a decrease in PV pH immediately preceding induced or natural egress. The pH drop was partly dependent on expression of PLP1 and was entirely absent in parasites lacking PLP1 and motility. We also identified formate-nitrite transporters (FNTs) as contributing to the acidification of the PV during egress, likely via co-transporting protons with the glycolytic products lactate and pyruvate. However, disruption of FNTs did not completely abrogate the release of lactate and pyruvate or eliminate the drop in PV pH, suggesting the involvement of other unidentified transporters or products that also contribute to PV acidification during egress.

## Results

### PV pH decreases immediately prior to induced and natural egress

Green fluorescent protein (GFP) has two excitation peaks at 410 nm and 470 nm. This property has been exploited to create, via amino acid substitutions, pH sensitive variants termed pHluorins (14). Ratiometric pHluorin (RatpH) responds to a decrease in pH with reduced fluorescence from excitation at 410 nm and increased fluorescence from excitation at 470 nm. To measure the pH of the *T. gondii* PV, we expressed a codon-optimized RatpH in RH parasites (RH-RatpH). We then tested the ability of RatpH to respond to changes in pH by incubating RH-RatpH infected cells in buffers of defined pH containing nigericin to equilibrate H^+^ across membranes (**Figure 1A**). Obtaining ratiometric images by excitation at 410 nm and 470 nm (**Figure 1B**) allowed the generation of a calibration curve for pH (**Figure 1C**). To initially quantify changes in pH during egress we performed live ratiometric imaging of RH-RatpH infected cells in Ca^2+^ containing buffer upon adding the Ca^2+^ ionophore ionomycin, which induces egress by equilibrating calcium across membranes. Collecting a series of ratio images following ionomycin treatment showed that a decrease in PV pH occurs prior to rupture of the PVM, which was indicated by the release of RatpH from the PV in non-ratioed images (**Figure 1D**). Tracings from individual PVs showed that in contrast to a consistent pH observed upon treatment with vehicle (DMSO) (**Figure 1E, a**), PV pH decreased 10-30 sec prior to PV rupture in infected cells treated with ionomycin (**Figure 1E, b,c,d**). Although the magnitude of the decrease was variable, all the PVs we observed showed a drop in PV pH prior to PV rupture. Analysis of 35 PVs recorded over multiple experiments showed a mean decrease of 0.35 pH units, corresponding to a ∼3-fold increase in [H^+^] within the PV. We also observed a similar decrease in PV pH upon inducing egress with zaprinast (**Figure 1F**), which activates parasite protein kinase G upstream of Ca^2+^ signaling (15).

**Figure 1.**
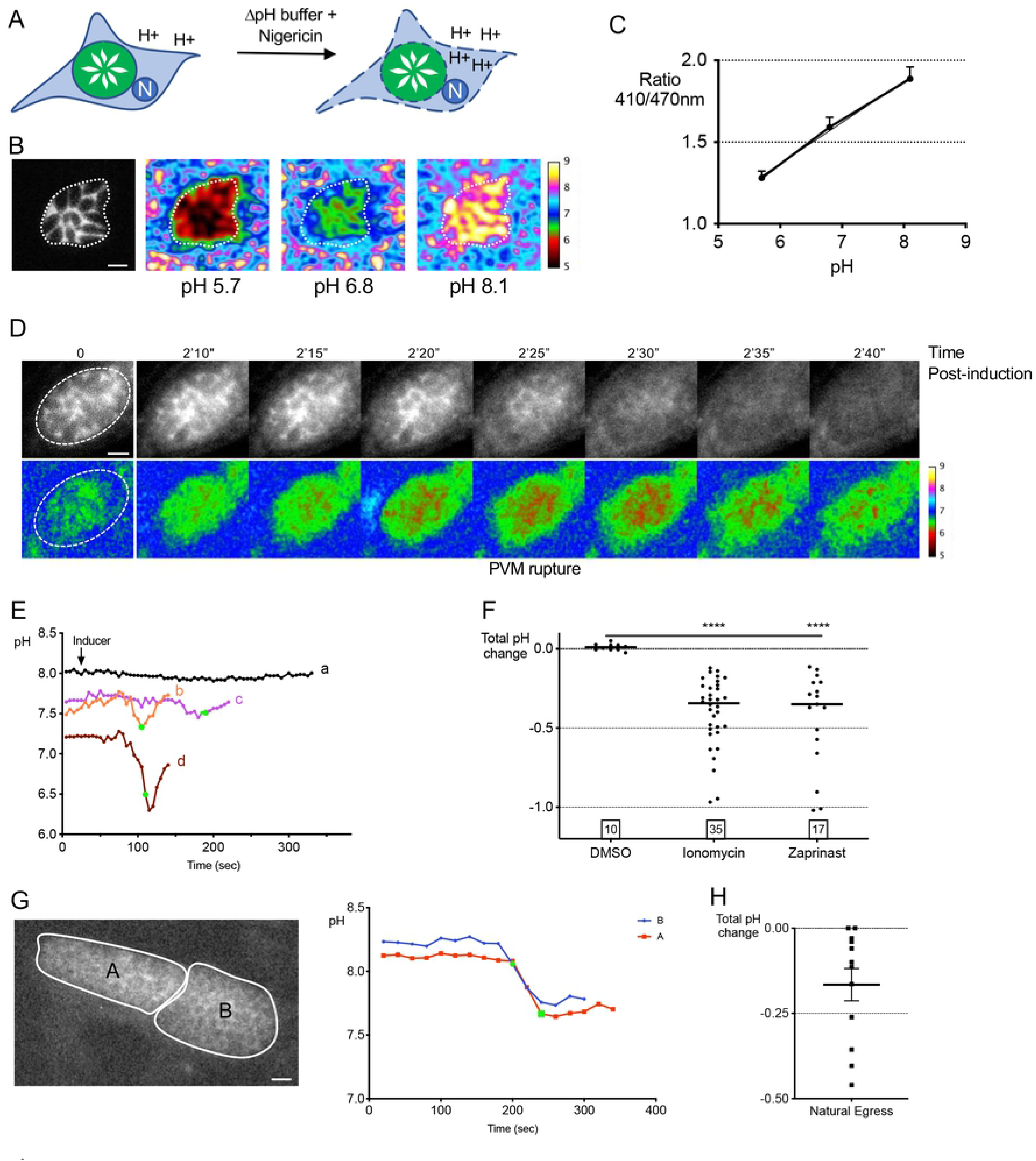
Measurement of pH in *T. gondii* PVs expressing ratiometric pHluorin. A) Equilibration of vacuoles to the surrounding buffer of known pH values with nigericin. B) Vacuole ratio images at 410/470 nm, and C) calibration curve from the ratio 410/470 nm values for each pH value (mean ± S.D); R^2^= 0.9992. Scale bar 5 μm. D) Reductions in PV pH during egress. Live imaging and time-course of 410/470 nm ratio images of induced RH-RatpH vacuole, pH scale bar indicated on the right. E) The magnitude of pH changes varies between PVs. Tracings of pH values of RH-RatpH PVs induced with DMSO (a) or ionomycin (b, c, d). Image acquisition was paused following frame 5 (25 sec), inducer was added, and acquisition was started again. Green data points indicate the time point pHuorin first leaves the PV. F) A significant reduction in pH in both ionomycin or zaprinast induced vacuoles. Data points represent changes in PV pH starting from baseline to a drop greater than 0.05 following induction with either ionomycin or zaprinast. Numbers in squares within the graph indicates the numbers of PVs enumerated. A Kruskal-Wallis test with Dunn’s multiple comparison was performed. **** *p*≤0.0001. Bars indicate the median. G) A drop in PV pH occurs in during natural egress. Representative PVs late in replication (∼50 h) and pH tracings associated with egress. Scale bar 10 μM. H) Quantification of pH changes in PVs during natural egress. Bar indicates the mean.

To determine if a decrease in pH occurs during natural egress, we imaged RH-RatpH infected cells at ∼48-52 h post-infection for 20 min/field of view. We observed a decrease in PV pH in 10 out of 12 egress events, with a mean decrease of 0.17 pH units (**Figure 1G, H**). Taken together, our findings suggest that PV pH decreases immediately prior to induced and natural egress. Due to the challenges of capturing natural egress events, all subsequent experiments were performed with induced egress.

### Ionic environment modulates the vacuolar pH response

Earlier work suggested that a loss of cytoplasmic K^+^ from the host cell triggers egress of *T. gondii* tachyzoites through activation of phospholipase C and a subsequent increase in cytoplasmic Ca^2+^ (16). A more recent study proposed that the decrease in K^+^ accelerates egress but is not a trigger (17). While Ca^2+^ is necessary for activation of egress, the absence of extracellular Ca^2+^ delayed but did not inhibit parasite egress (2). pH measurements presented thus far have been obtained in Ringer’s buffer (155 mM NaCl, 3 mM KCl, 1 mM Ca^2+^, 1 mM MgCl_2_, 10 mM glucose, 10 mM HEPES, 3 mM NaH_2_PO_4_, pH 7.4). To test the effect of Ca^2+^ and K^+^ in the extracellular media on the vacuolar pH drop, infected cells were incubated in Ringer’s or 145 mM K^+^ (termed high K^+^) to mimic the [K^+^] of the host cell cytosol, each with and without Ca^2+^. Buffer compositions used in this study are listed in **Table 1**.

**Table 1.** Ion composition of buffers used in this study.

|  | **Ringer's** | **Ringer's-Ca<sup>2+</sup>** | **High K<sup>+</sup>** | **High K<sup>+</sup>-Ca<sup>2+</sup>** | **EC-Na<sup>+</sup>** | **EC-Cl<sup>-</sup>** |
| --- | --- | --- | --- | --- | --- | --- |
| NaCl | 155 mM | 155 mM | 5 mM | 5 mM |  |  |
| KCl | 3 mM | 3 mM | 145 mM | 145 mM | 5.8 mM |  |
| CaCl<sub>2</sub> | 2 mM |  | 2 mM |  | 1 mM |  |
| MgCl<sub>2</sub> | 1 mM | 1 mM | 1 mM | 1 mM | 1 mM |  |
| Glucose | 10 mM | 10 mM | 10 mM | 10 mM | 10 mM |  |
| HEPES | 10 mM | 10 mM | 15 mM | 15 mM | 10 mM |  |
| NaH<sub>2</sub>PO<sub>4</sub> | 3 mM | 3 mM |  |  |  |  |
| EGTA |  | 1 mM |  | 1 mM |  |  |
| Choline-Cl |  |  |  |  | 142 mM |  |
| Na<sup>+</sup> glutamate |  |  |  |  |  | 155 mM |
| K<sup>+</sup> glutamate |  |  |  |  |  | 3 mM |
| Ca<sup>2+</sup> glutamate |  |  |  |  |  | 2 mM |
| MgSO<sub>4</sub> |  |  |  |  |  | 1 mM |

Egress from infected cells in Ringer’s buffer was induced with zaprinast (**Figure 2A**) or ionomycin (**Figure 2B**) with or without cytochalasin D (CytD) treatment to determine whether immobilizing the parasites affects the decrease in PV pH after induction. Since CytD did not markedly affect PV pH, to facilitate imaging it was included in all subsequent experiments unless otherwise noted. Whereas removal of Ca^2+^ from Ringer’s significantly attenuated the drop in PV pH after induction with zaprinast or ionomycin, removal of Ca^2+^ from the high K^+^ had no effect. In the presence of Ca^2+^, incubation in high K^+^ showed a trend toward attenuating the drop in PV pH after induction with zaprinast and a significant attenuation following ionomycin induction. When Na^+^ in the extracellular buffer is replaced with choline to maintain the total concentration of monovalent cations, with or without Ca^2+^, there was no effect on the pH drop observed (**Figure 2C**). A role for chloride was assessed by replacing it with glutamate or sulfate (-Chloride buffer) composed of potassium glutamate, sodium glutamate, calcium glutamate, and magnesium sulfate. The absence of chloride also had no effect on PV acidification (**Figure 2C**). Together these findings imply that extracellular Ca^2+^ is necessary for normal acidification of the PV during egress and that the loss of K^+^ from host cells also influences PV pH.

**Figure 2.**
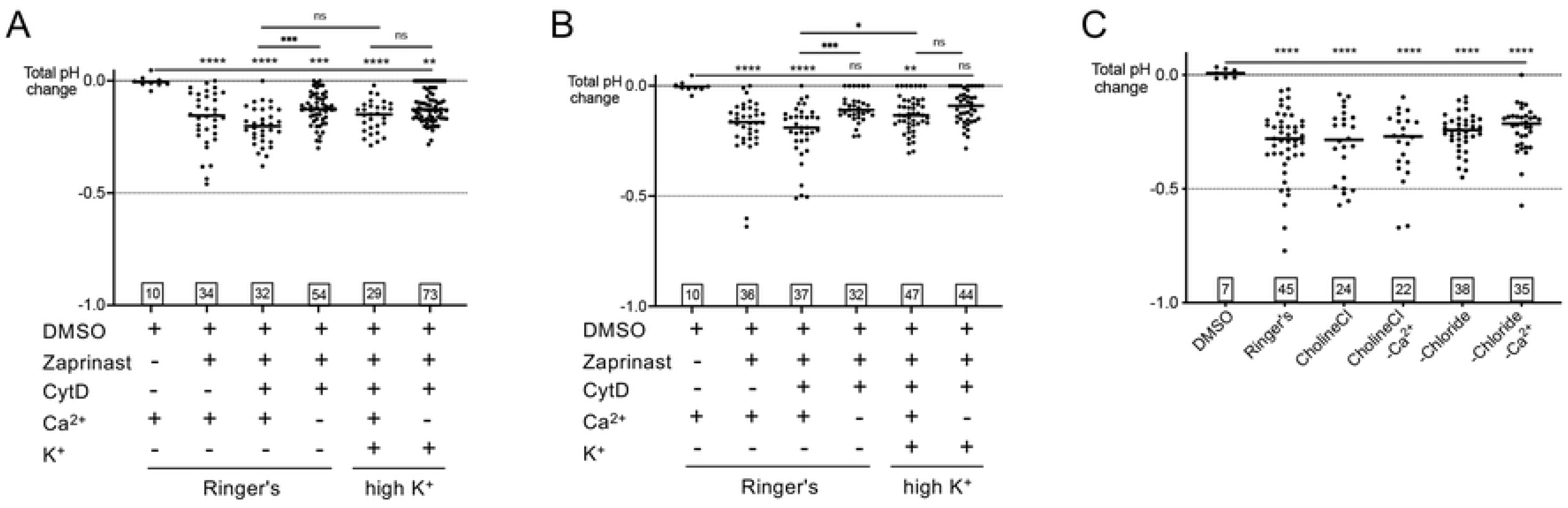
Composition of surrounding buffer mildly abates PV pH changes. A) and B) Extracellular buffer without Ca^2+^ tempers the drop in pH with ionomycin (A) or zaprinast (B) induction. RH-RatpH PVs, treated with or without cytochalasin D, incubated in Ringer’s buffer or High K^+^ with or without Ca^2+^ and induced with DMSO or zaprinast. C) Na^+^ and Cl^-^ do not affect pH changes. Extracellular buffer with Na^+^ replaced with choline or all Cl^-^ replaced with glutamate, with or without Ca^2+^. Numbers in squares within all graphs indicate the numbers of vacuoles enumerated. A Kruskal-Wallis test with Dunn’s multiple comparison was performed for all graphs. Bars indicate the mean.

### PLP1 influences acidification of the PV

Parasites deficient in *PLP1* were previously shown to be substantially delayed in induced egress, with a proportion of the parasites unable to leave the vacuole (18). Since PLP1 is a pore-forming protein, we reasoned that it could facilitate ion flux associated with acidification of the PV during egress. To address this, we introduced RatpH into PLP1 deficient parasites (Δ*plp1*-RatpH). As expected, Δ*plp1*-RatpH showed a delay in egress, with parasite egress occurring substantially later than the onset of motility (**Figure 3A**). We found that although Δ*plp1*-RatpH vacuoles from which the parasite was able to egress exhibited a drop in pH with ionomycin or zaprinast induction, this decrease was significantly attenuated relative to RH-RatpH (**Figure 3B**). Whereas the measurements thus far were performed on the entire PV, we noted that some regions of the PV show greater acidification. Thus we also measured the minimum pH in each PV prior to egress. This analysis showed that the mean difference in the lowest regional PV pH for non-induced (DMSO) and induced (zaprinast or ionomycin) RH-RatpH parasites is ∼0.7 pH units (**Figure 3C**). The analysis of lowest regional pH also confirmed an attenuation of PV acidification during induced egress of Δ*plp1*-RatpH parasites.

**Figure 3.**
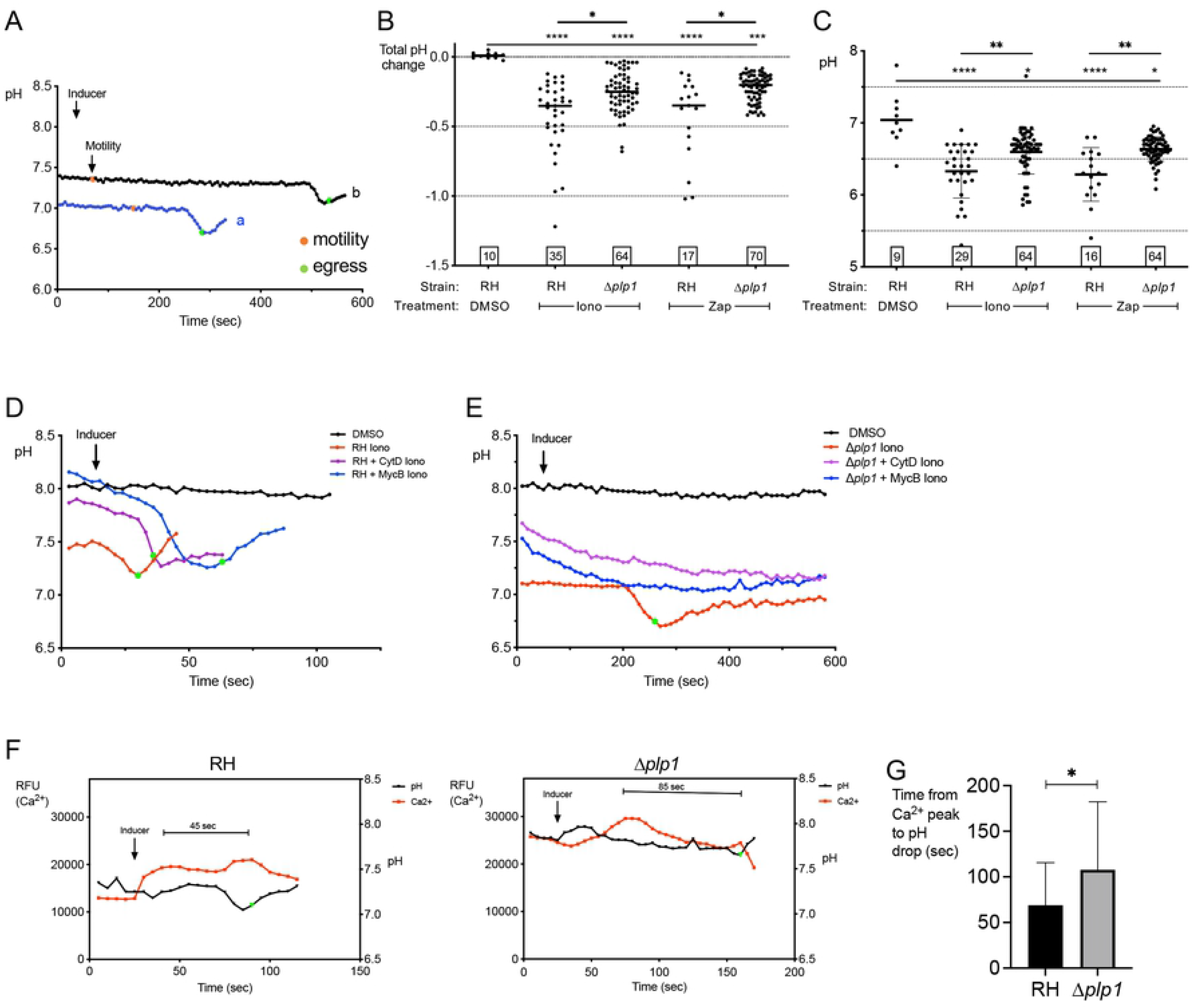
PV acidification occurs in Δ*plp1* parasites. A) Tracings of pH values in Δ*plp1*-RatpH from live imaging. The orange data point indicates the time point where motility of the tachyzoites began and the green points indicate the time point that pHluorin leaves the vacuole. Two representative tracings are shown. Arrow indicates the time point that inducer was added. B) Δ*plp1* PVs show a significant attenuation of PV acidification after induction with ionomycin or zaprinast. RH-RatpH data sets are the same as those presented in Figure 1F. Data points represent PV changes in pH starting from baseline to a drop greater than 0.05 following induction with either ionomycin or zaprinast. Numbers in squares within the graph indicates the numbers of PVs enumerated. A Kruskal-Wallis test with Dunn’s multiple comparison was performed. * *p*≤0.05, ** *p*≤0.01, **** *p*≤0.001, **** *p*≤0.0001. C) Punctate foci in individual vacuoles reach lower pH than whole vacuole. Ratio images of RH-RatpH or Δ*plp1*-RatpH vacuoles were analyzed in ImageJ using the Process Plugin to find min-max values (v.1.00). Data points indicate the lowest pH measured within a PV. Bars indicate the mean. D) Paralyzed WT parasites undergo PV acidification. Representative traces of RH-RatpH parasites untreated or immobilized with cytochalasin D (CytD) or mycalolide B (MycB), followed by ionomycin induction. E) Representative traces of Δ*plp1*-RatpH parasites untreated or immobilized with CytD or MycB, followed by ionomycin induction. F) A Ca^2+^ increase follows the addition of inducer and always precedes egress. Representative traces of relative Ca^2+^ levels (left axis) and pH (right axis) of RH-RatpH (left graph) and Δ*plp1*-RatpH (middle graph) PVs following ionomycin induction. Enumeration of 41 RH-RatpH and 20 Δ*plp1*-RatpH PVs shows an increase in the time from Ca^2+^ increase to PV acidification in Δ*plp1*-RatpH (right graph). Error bars indicate the mean ± S.D.

While *PLP1*-deficient parasites are defective in egress, some Δ*plp1* parasites are capable of egressing, most likely due to the motility of the parasites that eventually break through the PVM and plasma membrane of host cells. To more definitively study the role of PLP1 and motility in the pH drop associated with egress, RH-RatpH or Δ*plp1*-RatpH parasites were paralyzed with either cytD or mycalolide B (MycB), which prevent actin polymerization or severs F-actin, respectively. As observed earlier in this study (Figure 2A and 2B), PVs of immobilized RH-RatpH parasites induced with ionomycin still displayed a significant drop in pH associated with pHluorin release (**Figure 3D**). However, PVs of immobilized Δ*plp1*-RatpH showed no decrease in pH and they failed to release pHluorin for the entire duration of the experiment (up to 20 min) in all 225 vacuoles observed (**Figure 3E**). Taken together, these findings suggest that PLP1 influences PV pH and that rupture of the PVM and possibly the host plasma membrane via PLP1 or motility is necessary for acidification of the PV.

The lack of vacuolar acidification of PLP1-deficient and immobilized tachyzoites suggests that parasite factors initiate and are necessary for the pH drop. To explore whether acidification could occur first in the host cell and then affect the PV pH, ratiometric pHluorin under a mammalian promoter was transfected into HeLa cells and infected with RH-RatpH (**Supplemental Figure S1A and S1B)**. Ratio images following induction with ionomycin in this example and others seem to indicate a pH change first occurring in the PV, followed by acidification in the host cell (**Supplemental Figure S1C)**. However, pH measurements of the PV and host cell show a close, indistinguishable pattern of the pH changes (blue and green tracings in **Supplemental Figure S1D),** and a clear drop occurring first in either the PV or host cannot be distinguished. Nevertheless, the combined results of the absence of a pH change in immobilized Δ*plp1*, and that the pH of the host cell does not change in transfected but uninfected host cells upon ionomycin or zaprinast induction (**Supplemental Figures S1D,E),** support the hypothesis that PV acidification is dictated by the parasite.

Since Ca^2+^ signaling triggers the secretion of microneme proteins including PLP1 and it activates motility, we sought to define the dynamics of parasite cytosolic Ca^2+^ and their relationship to pH changes during egress by stably expressing a red genetically encoded Ca^2+^ indicator RGECO in the cytosol of RH-RatpH and Δ*plp1*-RatpH parasites. Ca^2+^ and PV pH measurements were obtained from non-immobilized parasites so that we could define the timing of Ca^2+^ and pH dynamics in relation to egress. RH-RatpH-RGECO parasites displayed a rapid increase in cytosolic Ca^2+^ after ionomycin induction and well before the drop in PV pH and egress (**Figure 3F** and **Supplemental Figure S2**). The decrease in PV pH often corresponded to a second peak of Ca^2+^, the presence of which has been reported previously (2). In contrast, Δ*plp1*-RatpH-RGECO parasites typically showed a single peak that developed more slowly, with a time interval being close to twice as long as that seen in RH-RatpH-RGECO parasites (**Figure 3G**). From these findings we conclude that Ca^2+^ signaling initiates prior to acidification of the PV and that the timing of Ca^2+^ signaling is delayed in parasites lacking PLP1.

### Several plasma membrane proton transporters are not required for acidification of the PV

Next, we sought to identify the basis for acidification of the PV by initially focusing on proton transporters with the potential to be expressed on the parasite surface. A V-type H^+^ ATPase (V-ATPase) is localized on the plasma membrane as well as to the plant-like vacuole (PLV)/Vacuolar compartment (VAC, used hereafter) organelle. The V-ATPase was shown to function in maintaining cytoplasmic pH and the acidic pH of the VAC and immature rhoptries (19). To test if the V-ATPase contributes to acidification of the PV, we transiently expressed RatpH in the inducible knockdown parasite strain of VHA1 (iVHA), a critical subunit of V-ATPase, generated in Stasic et al. We confirmed down-regulation of VHA1 protein levels following ATc treatment by western blotting for the HA tag appended to VHA1 (**Figure 4A**). We found that parasites lacking VHA1 (iΔ*vha1*-HA +ATc) showed a drop in PV pH after egress induction that was indistinguishable from those expressing VHA1 (iΔ*vha1*-HA) (**Figure 4B,C**). These findings suggest that VHA1 does not play a role in PV acidification during egress.

**Figure 4.**
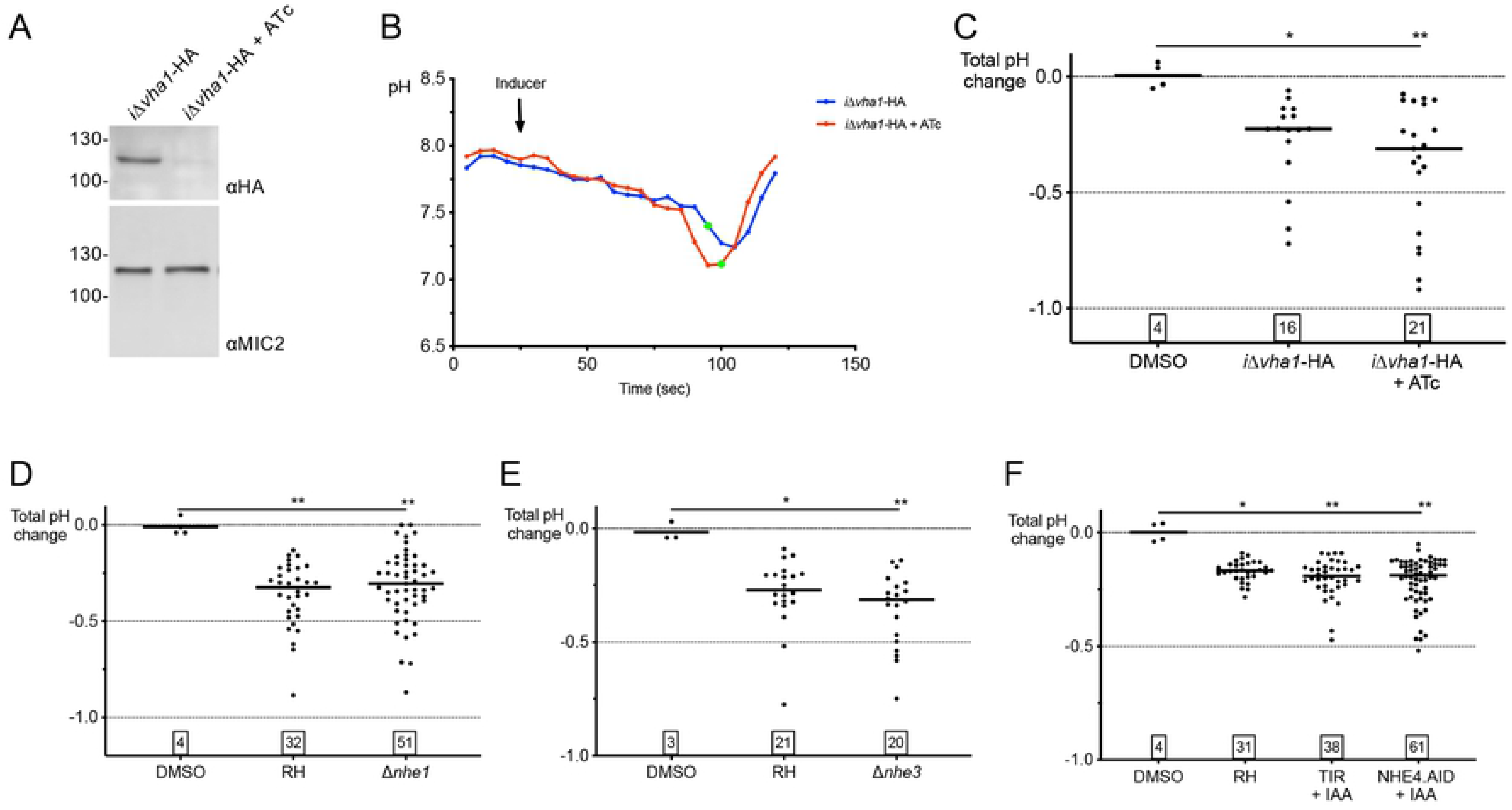
Known plasma membrane-localized V-ATPase and sodium-hydrogen exchangers do not affect vacuolar pH changes during egress. A) Western blot of inducible VHA1 (*iΔvha1*-HA) +/- anhydrotetracycline (ATc) treatment for 32h. Blots were probed with Ms anti-HA to detect VHA1-HA; Rb anti-MIC2 was a loading control. B) Representative pH tracings of *iΔvha1*-HA parasites transiently transfected with RatpH, incubated +/– ATc, and induced with ionomycin. The green data points indicate the time point that pHluorin leaves the vacuole. C) Knockdown of *iΔvha1*-HA does not affect the magnitude of a pH change. D-F) Sodium-proton exchangers do not contribute to vacuolar pH changes. Parasite strains Δ*nhe1* (D), Δ*nhe3* (E), NHE4.AID + IAA (F) transiently transfected with RatpH were induced with ionomycin and pH changes measured by live-imaging. Numbers in squares within graphs indicate the numbers of vacuoles enumerated. Kruskal-Wallis tests with Dunn’s multiple comparison was performed for all graphs. * *p*≤0.05, ** *p*≤0.01. Bars indicate the median.

*T. gondii* expresses four sodium/proton exchangers: NHE1 is on the plasma membrane (Arrizabalaga et al., 2004), NHE2 is associated with the rhoptries (22), NHE3 is associated with the VAC (23), and NHE4 is predicted to be on the plasma membrane or Golgi, depending on the prediction program used for the analysis (Barylyuk et al., 2019)(20, 21). Since it was unlikely that a rhoptry proton exchanger would affect PV acidification during egress, NHE2 was excluded from testing. We tested available knockout strains that lack NHE1 (20) or NHE3, (23); which we confirmed by PCR (**Supplemental Figure S3**). Upon transiently expressing RatpH, we found that Δ*nhe1* and Δ*nhe3* both showed normal acidification of the PV during induced egress (**Figure 4D,E**). To assess the potential contribution of NHE4 to pH changes in the PV during egress, we used CRISPR-Cas9 to append an auxin-inducible degron (AID) to the C-terminus of NHE4 (NHE4.AID) in RHΔ*ku80* parasites expressing the auxin receptor (TIR1) for AID-based protein degradation (RHΔ*ku80*/TIR, TIR hereafter) (24). Immunofluorescence localization of the HA tag downstream of the AID indicated that the tagged NHE4 was found primarily in the region anterior to the nucleus, reminiscent of Golgi staining (**Supplemental Figure S4A**). Addition of auxin (indoleacetic acid, IAA) effectively reduced expression of NHE4.AID to a level below detection based on western blotting (**Supplemental Figure S4B**). NHE4.AID parasites showed an acidification of the PV that was indistinguishable from that of RH or TIR parasites (**Figure 4F**). Altogether, this showed that NHE1, NHE3, and NHE4 do not contribute to the vacuolar acidification observed during egress.

### FNT1 and FNT2 are important for acidification of the PV during egress

Formate-nitrite transporters (FNTs) are multi-pass transmembrane proteins that transport monocarboxylate metabolites, such as formate and lactate, in *P. falciparum* (25, 26) and in *T. gondii* (27). *T. gondii* expresses three FNTs (FNT1-3) that localize to the parasite plasma membrane upon overexpression in tachyzoites (27). Proteomics and transcriptomics data available in Toxodb suggest that FNT1 and FNT2 proteins are primarily expressed in tachyzoites whereas FNT3 is mainly expressed in the feline enteroepithelial stages. A previous study determined that TgFNTs transport L-lactate and formate in a pH-dependent manner, and it described small molecule inhibitors of FNT1-3 that impaired the tachyzoite lytic cycle (27). More recent gene knockout studies reported that FNT1 plays a prominent role in lactate export (28, 29).

To initially assess whether FNTs had a role in PV acidification, the two most potent inhibitors of parasite growth from the Erler et al study, BH-296 and BH-388 (27), were incubated with infected cells and PV pH changes during time-lapse microscopy were determined in RH-RatpH vacuoles. We initially tested the concentrations of the inhibitors that were used in the previous study, but 10 μM, BH-296 blocked tachyzoite egress, precluding its use at this concentration. At both 1 μM and 5 μM, BH-296 reduced the magnitude of pH change during zaprinast-induced egress compared to RH-RatpH, whereas BH-388 only had an affect at 10 μM (**Figure 5A**).

**Figure 5.**
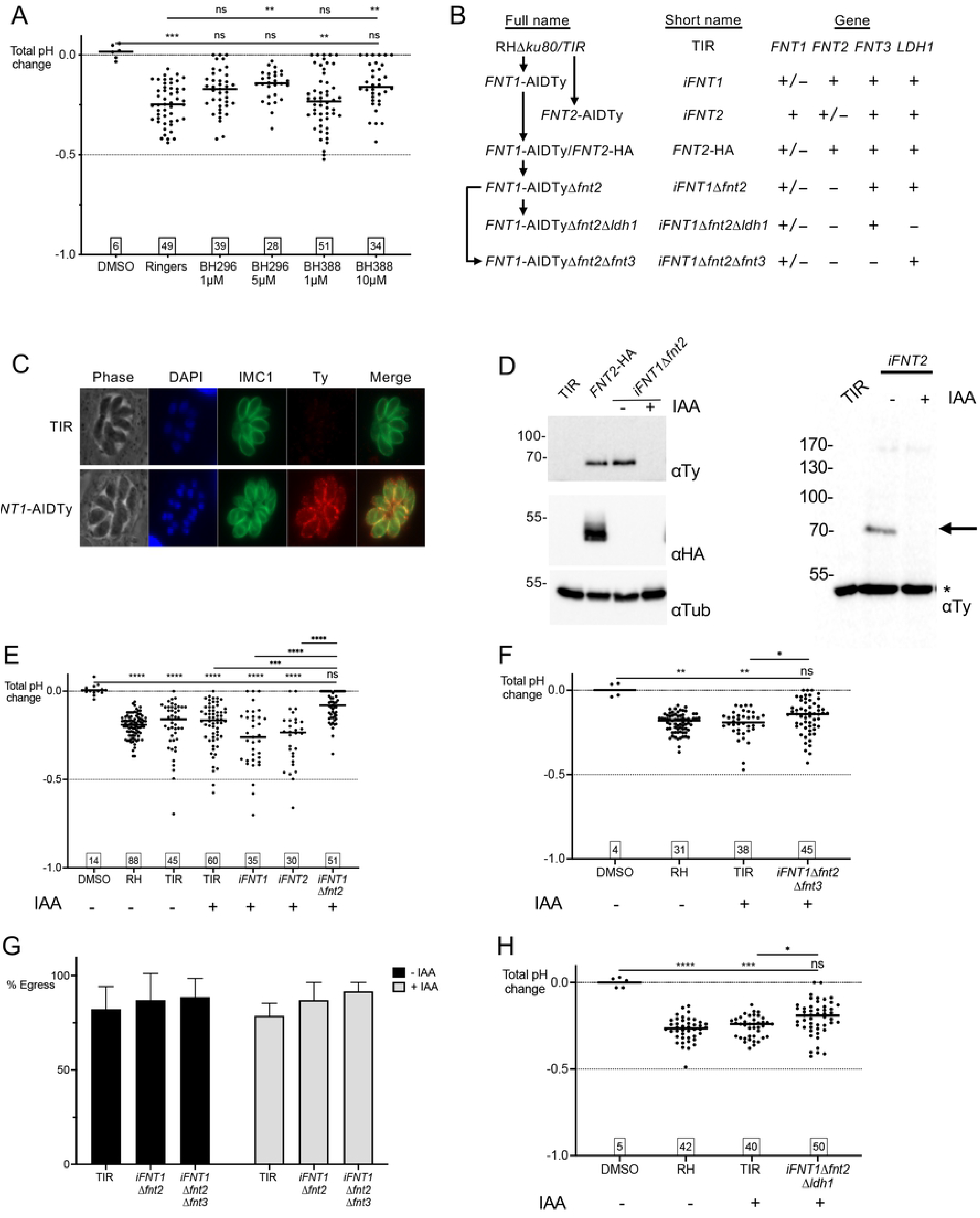
Impairing formate-nitrite transporters attenuates PV acidification. A) Effect of FNT inhibitors BH-296 and BH-388 at 1 μM, 5 μM, or 10 μM on pH changes during zaprinast-induced egress. B) Schematic description of FNT strains generated, with full and short names used in graphs. C) Immunofluorescence staining of parental TIR and FNT1-AIDTy intracellular vacuoles with anti-IMC1 and anti-Ty antibodies. D) IAA-treatment down-regulates FNT1 and FNT2 expression. Parasites treated for 24 h with or without IAA were purified and lysates were immunoblotted with anti-Ty; anti-HA showed expression of FNT2-HA. * indicates a non-specific band recognized by the anti-Ty Ab. E) Disruption of FNT1 and FNT2 reduces the magnitude of PV acidification. F) Knockout of *FNT3* does not further attenuate the pH drop. G) Static egress assays of 28 h vacuoles of parental TIR and i*FNT1*Δ*fnt2*Δ*fnt3* ± IAA treatment. Error bars indicate the mean ± S.E.M. At least 250 PVs were enumerated. H) Knockout of *LDH1* does not further attenuate the pH drop. Numbers in squares within graphs indicate the numbers of vacuoles enumerated. Kruskal-Wallis tests with Dunn’s multiple comparison was performed for all graphs. * *p*≤0.05, ** *p*≤0.01, **** *p*≤0.001, **** *p*≤0.0001. Bars indicate the median.

Since the inhibitors could target one or all the FNTs in *T. gondii* as well as potentially in the host cell, we created transgenic parasites to identify the contribution of FNT1 and FNT2 individually and together to the pH drop during egress; FNT3 was not initially examined given its lack of expression in the tachyzoite stage. A schematic of the strategy and the strains generated is illustrated in **Figure 5B** and **Supplemental Figure S5A**. The mini auxin-induced degron (mAID) cassette, including an mNeonGreen (mNG) and a Ty epitope tag (mNG-AIDTy), was introduced into the C-terminus of FNT1 by CRISPR-Cas9. Live imaging of the mNG tag as well as immunofluorescence assays of the Ty tag indicated the proper targeting and localization of the cassette in FNT1 (**Figure 5C** and **Supplemental Figure S6A**) and correct tagging was confirmed by PCR (**Supplemental Figure S5B**) and western blot (**Figure 5D**). The mNG-AIDTy cassette was also introduced into the C-terminus of FNT2 (*iFNT2*) and confirmed by western blot (**Figure 5D**) but the mNG signal could not be detected by live imaging. The FNT1-AIDTy strain (*iFNT1*) was then used as the background in which FNT2 was tagged at the C-terminus with a 6xHA tag and HXGPRT selectable marker, generating the strain *FNT1*-AIDTy/*FNT2*-HA (*FNT2*-HA). Detection of the 6xHA could not be observed by immunofluorescence, similar to the inability to detect the Ty tag in *iFNT2*, perhaps due to low expression of FNT2. However, PCR (**Supplemental Figure S5B**) and western blotting (**Figure 5D**) confirmed the integration and expression of the HA tag, respectively. The *FNT2* coding region was deleted using gRNAs targeting the 5’ end of *FNT2* and the 3’ end of the HXGPRT selectable cassette by replacement with the DHFR-TS selectable marker, generating the *iFNT1*Δ*fnt2* strain (**Supplemental Figure S5A**). Integration of the DHFR-TS in the FNT2 locus was confirmed by PCR (**Supplemental Figure S5B**). The effectiveness of FNT1 knockdown with IAA was determined by both live imaging of the mNG tag (**Supplemental Figure S6A**) and by western blotting for the Ty tag (**Figure 5D** and **Supplemental Figure S6B**). FNT1-AIDTy was essentially undetectable after 6 h of IAA treatment. Transient transfection of *iFNT1* parasites with RatpH and incubation with IAA for 20 h was performed prior to time-lapse imaging. We found that although inducible knockdown of FNT1 (*iFNT1* + IAA) or FNT2 (*iFNT2* + IAA) did not affect PV pH during egress, inducible knockdown of FNT1 in the absence of FNT2 (*iFNT1Δfnt2* + IAA) significantly attenuated the decrease in PV pH during egress, albeit incompletely. The PVs of some *iFNT1Δfnt2* + IAA parasites showed no detectable decrease in PV pH during egress, suggesting that PV acidification is not critical for parasite exit from host cells. *iFNT1* and *iFNT2* parasites without IAA treatment could not be analyzed by live imaging due to spectral overlap of mNG with RatpH. These findings suggest that FNT1 and FNT2 are involved in the observed pH drop in the PV during induced egress. Additionally, the magnitude of the pH drop in these strains being comparable to those observed for the inhibitors is supportive of the inhibitors targeting the parasites specifically versus the host FNTs.

### FNT3 does not play a role in PV acidification

Although FNT3 appears to be expressed almost exclusively in the sporozoite stages of the parasite based on Toxodb, compensation of FNT1 and FNT2 transport by upregulation of FNT3 expression in the absence of FNT1 and FNT2 was a possibility. To measure potential changes in FNT3 expression by reverse transcriptase quantitative PCR, we tested two independent primer sets to the FNT3 mRNA and used actin as the housekeeping gene for normalization. No significant change in FNT3 transcript level was observed in the absence of FNT1 and FNT2 (**Supplemental Figure S7**). To more conclusively rule out a role for FNT3 for the residual drop in PV pH observed in *iFNT1*Δ*fnt2*, we deleted FNT3 to generate an *iFNT1*Δ*fnt2*Δ*fnt3* strain (**Supplemental Figure S5A**). We found that deletion of FNT3 in the absence of FNT1 and FNT2 did not further exacerbate the attenuation of PV acidification during egress, and that a residual drop of PV pH is still observed (**Figure 5F**). Despite the attenuation of a pH change in the PV during egress, neither the *iFNT1*Δ*fnt2* nor the *iFNT1*Δ*fnt2*Δ*fnt3* parasites were defective in their ability to egress upon induction with zaprinast in a static (single 2 min time point) egress assay (**Figure 5G**). The absence of these FNTs also had no effect on intracellular replication (**Supplemental Figure S8**). Taken together our findings suggest that FNT1 and FNT2 contribute to acidification of the PV during egress, acidification is not critical for induced egress, and other transporters likely exist for residual acidification of the PV in parasites lacking the FNTs.

### Release of Lactic acid and pyruvic acid potentially contribute to acidification of the PV

L-lactate is a major by-product of glycolysis. FNT1 was recently shown to be the principal lactate transporter in *T. gondii* tachyzoites (29). Lactate dehydrogenases (LDHs) convert the end-product of glycolysis, pyruvate, to lactate. *T. gondii* harbors two LDHs, LDH1 and LDH2, with LDH1 being predominantly expressed in tachyzoites and LDH2 being expressed exclusively in the bradyzoite stage (30, 31). Pomel and colleagues (32) reported that several glycolytic enzymes including LDH1 relocalize from the cytosol to the parasite periphery during the transition from intracellular to extracellular parasites. The authors suggested this relocation could underlie subpellicular production of ATP from glycolysis for actinomyosin-dependent gliding motility during and following egress. However, several studies have reported that although Δ*ldh1* parasites are virulence attenuated and have defects during the chronic stage, they are not defective in the lytic cycle *in vitro* (30,33,34). To determine whether the generation of lactate by LDH1 contributed to the residual pH drop observed in the *iFNT1*Δ*fnt2* strain, we disrupted *LDH1* by replacement with the bleomycin resistance cassette (**Supplemental Figure S5A**). The resulting *iFNT1*Δ*fnt2Δldh1* strain was transiently transfected with RatpH and pH changes during induced egress were measured. The loss of LDH1 in addition to FNT1 and FNT2 did not further attenuate the pH drop beyond that observed for *iFNT1*Δ*fnt2* alone (**Figure 5E,H**), indicating that, while lactate release into the PV could contribute to the pH drop during egress, other molecules clearly play a role in this function.

The findings from *iFNT1*Δ*fnt2Δldh1* parasites imply that lactate production is not essential for the observed pH drop during egress. To determine whether the release of lactate from the parasites could contribute to the changes in PV pH, we simulated the release of products during egress from TIR and FNT transgenic strains (pretreated with or without IAA) by collecting excreted-secreted antigen (ESA) supernatants from extracellular parasites induced with zaprinast or solvent control. ESAs were then assayed for lactate and pyruvate. For the TIR parental strain, we found that zaprinast treatment showed a trend toward increasing the release of lactate, but the difference was not statistically significant (**Figure 6A**). Unexpectedly, *iFNT1* and *iFNT2* without IAA treatment both showed a significant reduction in lactate release relative to the parental strain (**Figure 6B**), implying that the mNG-AIDTy tag interferes with the function of these two transporters. That these strains showed a significant reduction in lactate release without an effect on PV pH indicates either that the level of lactate released is sufficient for the observed pH changes or other molecules are involved. The observed effects in *iFNT1* and *iFNT2* are inconsistent with the Zeng et al study (29), which suggested that FNT2 does not contribute significantly to the transport of L-lactate across the plasma membrane. Treatment with IAA did not further diminish the release of lactate, possibly due to the presence of the other transporter (FNT2 in *iFNT1* and FNT1 in *iFNT2*). Double (*iFNT1*Δ*fnt2*) and triple knockout (*iFNT1*Δ*fnt2*Δ*fnt3)* strains appear to release even less lactate, which is consistent with a role for FNT1 and FNT2 in acidification of the PV during egress. As expected, lactate release by *iFNT1*Δ*fnt2*Δ*ldh1* parasites was near the limits of detection. We conclude from these findings that release of lactate potentially contributes to the observed drop in PV pH during egress.

**Figure 6.**
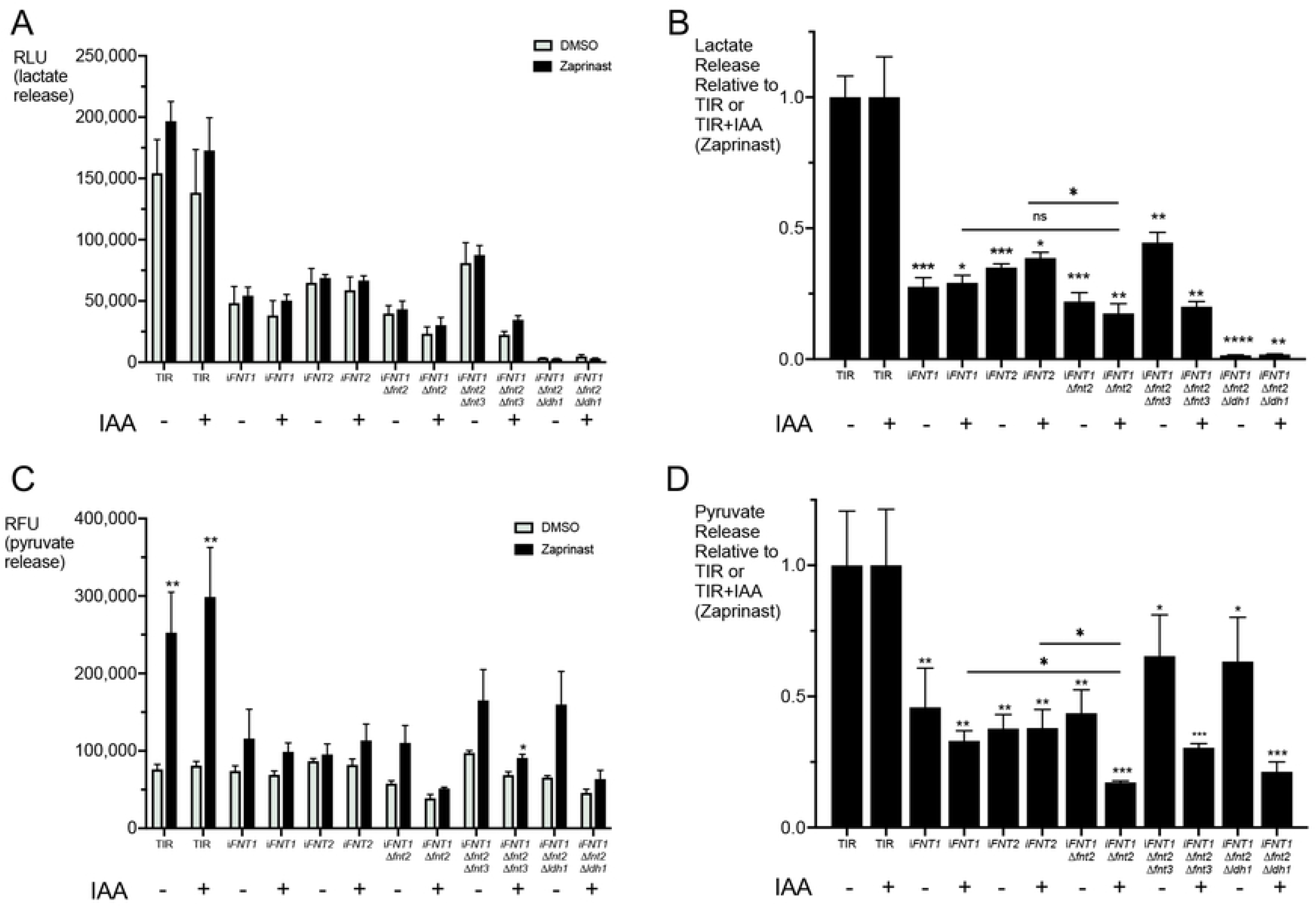
Release of lactate and pyruvate could contribute to PV pH changes during egress. A) Lactate release detected in excreted-secreted antigen (ESA) following DMSO (vehicle control) or zaprinast induction. RLU, relative luminescence units. B) Lactate release expressed relative to parental TIR parasites ± IAA. C) Pyruvate release detected in ESA following DMSO (vehicle control) or zaprinast induction. RFU, relative fluorescence units. D) Pyruvate release expressed relative to parental TIR parasites ± IAA. Two-tailed student’s *t*-test * *p*≤0.05, ** *p*≤0.01, **** *p*≤0.001, **** *p*≤0.0001. Error bars are mean ± S.E.M. Data in A) and B) represent 3 biological replicates with duplicates within each experiment. E) Proposed model of PV acidification during egress. An increase in cytosolic Ca^2+^ releases PLP1 into the PV to perforate the PVM and host cell. Lactate, pyruvate, and other ions or molecules are secreted into the PV through the FNTs and additional yet-to-be-identified transporters and channels to acidify the PV prior to egress. The drop in pH could further activate PLP1 in punctate sites within the PV for permeabilization.

A similar trend of pyruvate release was observed in the strains and treatments. Interestingly, we found that zaprinast treatment increased the release of pyruvate from the TIR parental strain by 2- to 3-fold (**Figure 6C**), suggesting a possible link between Ca^2+^ signaling and glycolysis. The relative release of pyruvate from TIR and FNT transgenic strains largely mirrored that of lactate (**Figure 6D**), consistent with the mNG-AIDTy tag impairing FNT1 and FNT2 transport function. Knocking down FNT1 in the absence of FNT2 (*iFNT1*Δ*fnt2*), FNT2 and FNT3 (*iFNT1*Δ*fnt2Δfnt3*) or FNT2 and LDH1 (*iFNT1*Δ*fnt2Δldh1*) resulted in a more pronounced decrease in pyruvate export than that of lactate, thus confirming a role for FNT1 in pyruvate export. We observed no difference in pyruvate release between *iFNT1*Δ*fnt2* and *iFNT1*Δ*fnt2Δfnt3*, as expected given the lack of FNT3 expression in tachyzoites. Parasites lacking all three FNTs (*iFNT1*Δ*fnt2Δfnt3*+IAA) still showed residual release of pyruvate, implying the existence of other unidentified transporters capable of exporting pyruvate. Taken together, our findings imply an association of Ca^2+^ signaling with glycolysis, roles for FNT1 and FNT2 in the export of lactate and pyruvate, and residual release of lactate and pyruvate by non-FNT transporters.

## Discussion

*T. gondii* egress from host cells is a tightly regulated and elaborate process with many upstream components, such as those that control microneme secretion including protein kinase G and Ca^2+^ -dependent protein kinases, downstream players in the gliding motility machinery, and cell-to-cell communication that coordinates the disassembly of the basal F-actin to disconnect parasites from one another (reviewed in (1). The parasite also secretes several proteins that contribute to egress including perforin-like protein 1 (*TgPLP1*), which is released from the micronemes (18). It was previously shown by Roiko et al (7) that TgPLP1 cytolytic and membrane binding activities are enhanced at low pH (5.4 to 6.9) compared to neutral pH (7.4). This study also confirmed an earlier report that low pH activates tachyzoite motility (6), and it extended insight by showing this occurs in part through the activation of microneme secretion, which is also necessary for motility. PLP1 activity and parasite motility both contribute to egress since the absence of one or the other delays egress but does not completely impair it (18, 35). The relationship between PLP1 activity and gliding motility at low pH was further linked when it was observed that a pH shift occurs during both induced egress and natural egress on a population scale (7). Our expression of ratiometric pHluorin in the PV allowed for the direct measurement of pH changes within an individual vacuole during induced egress. The data definitively showed that a drop in pH occurs during both induced and natural egress.

The extent to which PV acidification facilitates egress remains unclear. On one hand, the mean regional minimum PV pH after induction (∼6.3) is within the range of that for which PLP1 membrane binding and activity is enhanced (5.4-6.9) (7). Although the relationship between environmental pH and motility has not been defined extensively, parasite exposure to pH 5.4 (7) or 6.0 to 7.3 (6) is known to activate motility. These findings are consistent with PV acidification augmenting PLP1 activity and parasite motility. On the other hand, the decrease in pH prior to egress showed substantial variation between PVs, and in some cases parasites egressed from host cells without a discernible change in PV pH. Attempts to measure the effect of disrupting PV acidification using the weak base NH_4_Cl or the P-type ATPase inhibitor DCCD were ineffective in our live-imaging set-up, despite their reported efficacy in static egress assays (7). This discrepancy could be due to differences in the experimental set up or to the continuous exposure of the parasites to intense illumination during image acquisition, amongst other variables. Also, our inability to completely abrogate PV acidification by disrupting substrate transporters further limited defining the influence of PV pH on egress. Nevertheless, a drop in pH is not essential for exit from host cells, which is consistent with the event being regulated at multiple levels to ensure successful liberation.

The extracellular ionic environment has been proposed to affect the initiation and progression of egress, particularly the reduction of [K^+^] following the breakdown of the host plasma membrane (16) and the presence of environmental Ca^2+^ (2). We found that high extracellular (EC) [K^+^] or the absence of EC Ca^2+^ moderately attenuated PV acidification, suggesting that ion flux influences PV pH during egress. Recent studies measuring parasite cytosolic Ca^2+^ during egress reported that WT parasites typically show two peaks of Ca^2+^, with the second peak being dependent on influx of EC Ca^2+^ into the parasite from the environment (2, 17). Parasites lacking PLP1 only showed one Ca^2+^ peak after a considerable delay (17), which is consistent with our findings. We showed that PV acidification is not dependent on motility when PLP1 is present and that it is partially dependent on PLP1 when motility is functional; however, the absence of PLP1 and motility completely abrogates PV acidification and egress. We suggest that PLP1 functions to permeabilize the PV membrane and possibly the host plasma membrane ahead of egress, thereby resulting in ion fluxes including exposing the parasite to environmental Ca^2+^ to amplify Ca^2+^ signaling. This amplification serves to further enhance microneme secretion (including the release of more PLP1), motility, and acidification of the PV as a forward feedback mechanism to mediate exit from host cells. In this scheme, PLP1 is potentially both a facilitator and a benefactor of PV acidification, with these elements occurring in succession prior to and during egress, respectively.

Exchangers, transporters, and H^+^-ATPases on the parasite plasma membrane are plausible candidates to mediate the flux of H^+^ necessary for the observed decrease in PV pH. Despite the delayed egress defects observed in parasites lacking VHA1, NHE1, or NHE3 (19,20,23), we found that such mutants showed similar PV pH changes as parental or control parasites. Knockdown of an uncharacterized sodium/H+ exchanger, NHE4, also showed no effect on PV pH.

Several studies have identified formate-nitrite transporters (FNTs) as transporters of monocarboxylate metabolites, such as formate and lactate (27–29). FNT transport of substrates depends on the cotransport of H^+^, thus export of lactate or other substrates is intrinsically linked to acidification of the environment. Previous work suggested that FNT1 is the major FNT in tachyzoites (27–29). To determine the role of FNT1 and potentially FNT2 in acidification of the PV during egress, we endogenously tagged FNT1 and FNT2 with an mNG-AIDTy cassette. Individual knockdowns of FNT1 and FNT2 had no effect on the drop in pH, but lactate and pyruvate release assays suggest that the addition of the bulky mNG-AID tag on these transporters may have affected their functions. Another possible explanation is the previously reported auxin-independent degradation of AID-tagged proteins in a broad range of systems (36–38). Since antibodies to FNT1 or FNT2 are not available to detect endogenous protein levels, we cannot determine whether these tagged strains are downregulated in the absence of IAA. Regardless, taken together our findings suggest that FNT1 and FNT2 both contribute to the release of lactate and pyruvate (and cotransport of H^+^) during egress, but that other transporters can also export H^+^ for residual acidification of the PV in the absence of FNT1 and FNT2.

It was unexpected that the *iFNT2* parasites showed reduced lactate and pyruvate release given the recent finding that FNT2 contributes little to lactate transport in WT extracellular parasites (29). One potential explanation is that FNT2 plays a more prominent role in export of lactate, as tested in our study, versus uptake of lactate as examined in Zeng et al. A common finding between the two studies is that the lack of expression of all three FNTs had no effect on replication of *T. gondii* under *in vitro* culture conditions.

We did not pursue in great depth the link between Ca^2+^ signaling and activation of glycolysis because it was beyond the scope of the study. Nevertheless, we noted that elevation of parasite intracellular Ca^2+^ consistently preceded acidification of the PV, which aligns with a potential role for Ca^2+^ in activating glycolysis with concurrent excretion of lactate and pyruvate for acidification of the PV. Accordingly, our observation that zaprinast treatment markedly enhances export of pyruvate in an FNT dependent manner warrants further attention. That zaprinast augments release of pyruvate but not lactate implies that the fraction of lactate made under inducing conditions does not change but that increased production of pyruvate is managed by exporting it in a manner that is partly dependent on FNT1 and FNT2. In other systems Ca^2+^ is also known to activate mitochondrial production of ATP through oxidative phosphorylation (39). If the activation of Ca^2+^ signaling triggers ATP production from glycolysis and mitochondrial respiration, this presumably serves to meet the energy intensive demands of gliding motility. Confirming this would reveal an additional role for Ca^2+^ signaling during egress on top of its critical contributions to activating microneme secretion and motility. However, other explanations for zaprinast induced release of pyruvate are plausible. In addition to its action on PKG, zaprinast was also shown to inhibit mitochondrial pyruvate carrier activity, resulting in an accumulation and release of pyruvate and aspartate in mouse retina (Du et al., 2013). It is thus also possible that increased release of pyruvate from parasites is influenced by zaprinast inhibition of mitochondrial pyruvate carrier. We also cannot discount the possibility that zaprinast inhibition of host mitochondrial pyruvate carrier elevates pyruvate in infected host cells. Additional studies are required to determine the extent to which Ca^2+^ signaling influences glycolytic flux.

We present conclusive evidence for a drop in pH in the PV during both induced and natural egress, and we propose a model of the molecular mechanism by which this could occur (**Figure 7**). In WT parasites, an egress signal increases parasite cytosolic Ca^2+^ levels, leading to the secretion of PLP1 from the micronemes and permeabilization of the PVM, and an influx of Ca^2+^ from the host cell (or EC medium) further increases gliding motility and egress. The increase in Ca^2+^ could also trigger the net release of H^+^, coupled with lactate, pyruvate or other anions, to acidify the PV and create an environment more amenable to PLP1 cytolytic activity. FNT1 and FNT2 together contribute to the transport of the components for acidification, but clearly other transporters and exchangers can also function in this capacity. One potential candidate is the aquaporin water channel, AQ1, previously localized to the PLV/VAC compartment in tachyzoites (40). A second aquaporin (AQ2) is also present in the *T. gondii* genome (toxodb.org). Several aquaporins have been shown to transport lactate (reviewed in (41). A recent subcellular proteomics study suggested the localization of AQ1 in the Golgi or the plasma membrane (PM), depending on the prediction program used for the analysis (21). If a fraction of AQ1 (or AQ2) is localized to the PM, it could play a role in the export of lactate. Another possibility is that the transporters and exchangers tested in this study act in concert. Amiloride is a potent inhibitor of Na^+^/H^+^ exchangers (42, 43). RH-RatpH vacuoles treated with and without amiloride followed by induction with either zaprinast or ionomycin showed a significant abatement of a drop in pH in the PV (**Supplemental Figure S9).** Although disruption of NHE1, NHE3, and NHE4 individually did not affect PV acidification, it is possible that disruption of two or more exchangers could have a greater effect. Additionally, the presence of multiple putative transporters, exchangers, P-type ATPases and proton ATPases in the *T. gondii* genome implies a pool of potential contributors to the drop in pH during egress. Identifying additional players could be the focus of future work. While we have shown that a pH drop is not essential for parasites to leave the PV and host cell *in vitro*, this acidification could have a more critical role *in vivo* or in different stages of the parasite life cycle.

**Figure 7.**
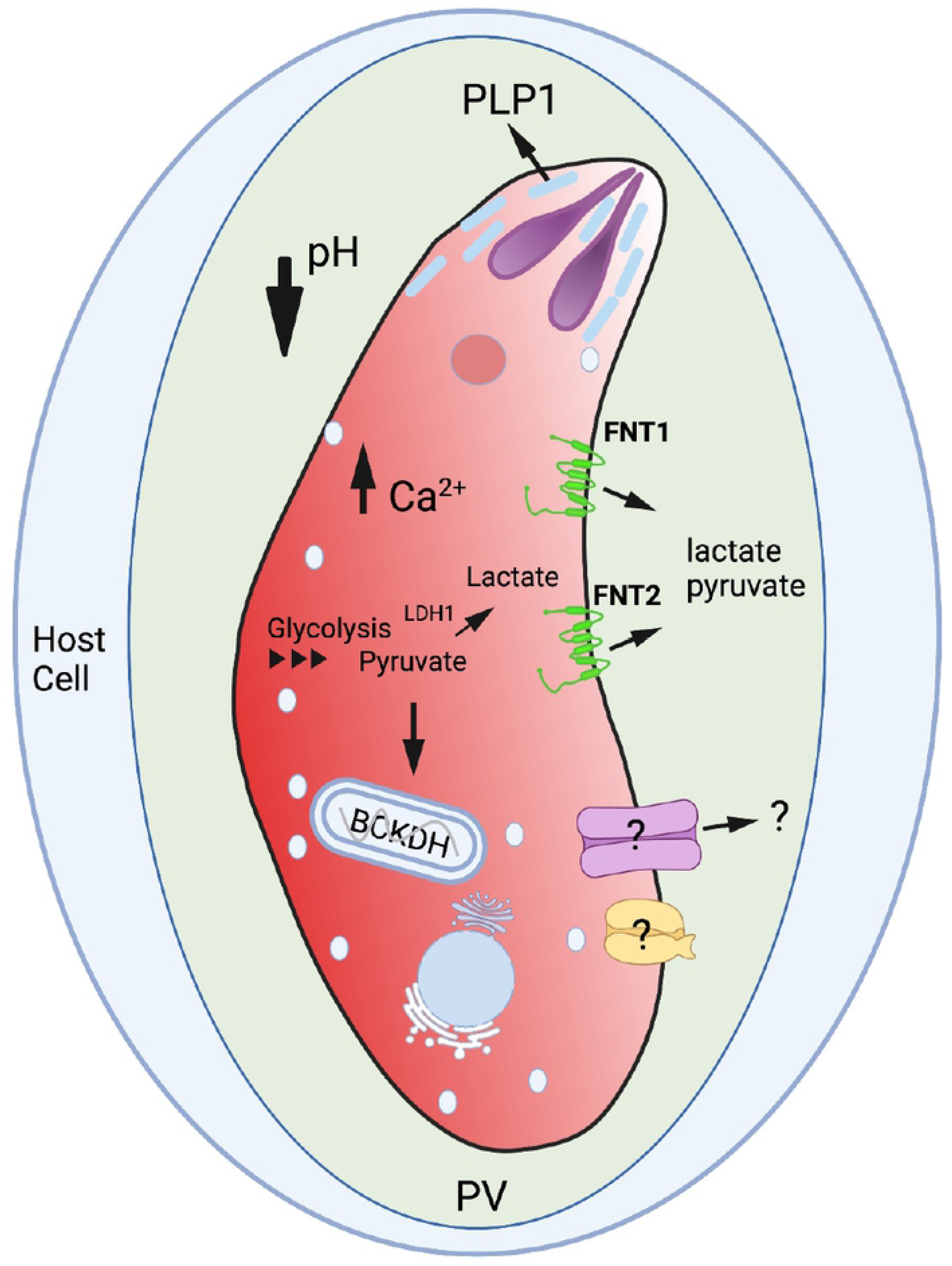
Model for pH acidification. Tachyzoites residing inside a parasitophorous vacuole (PV) within a host cell receive signal(s) that increase Ca^2+^ in the parasite, leading to secretion of PLP1 from the micronemes into the PV to form pores in the PV membrane and possibly host cell membrane. The products of glycolysis, lactate and pyruvate, are released into the PV space via formate-nitrite transporters (FNT1 and FNT2). These products, in addition to other molecules via other unidentified transporters, reduce the PV pH prior to egress. BCKDH, branched chain ketoacid dehydrogenase.

## Materials and methods

### Plasmids and parasite strains

Ratiometric pHluorin (RatpH) (14) was codon-optimized and chemically synthesized by GenScript Inc, as described previously (7). All primers used are found in **Table 2**.

**Table 2.**
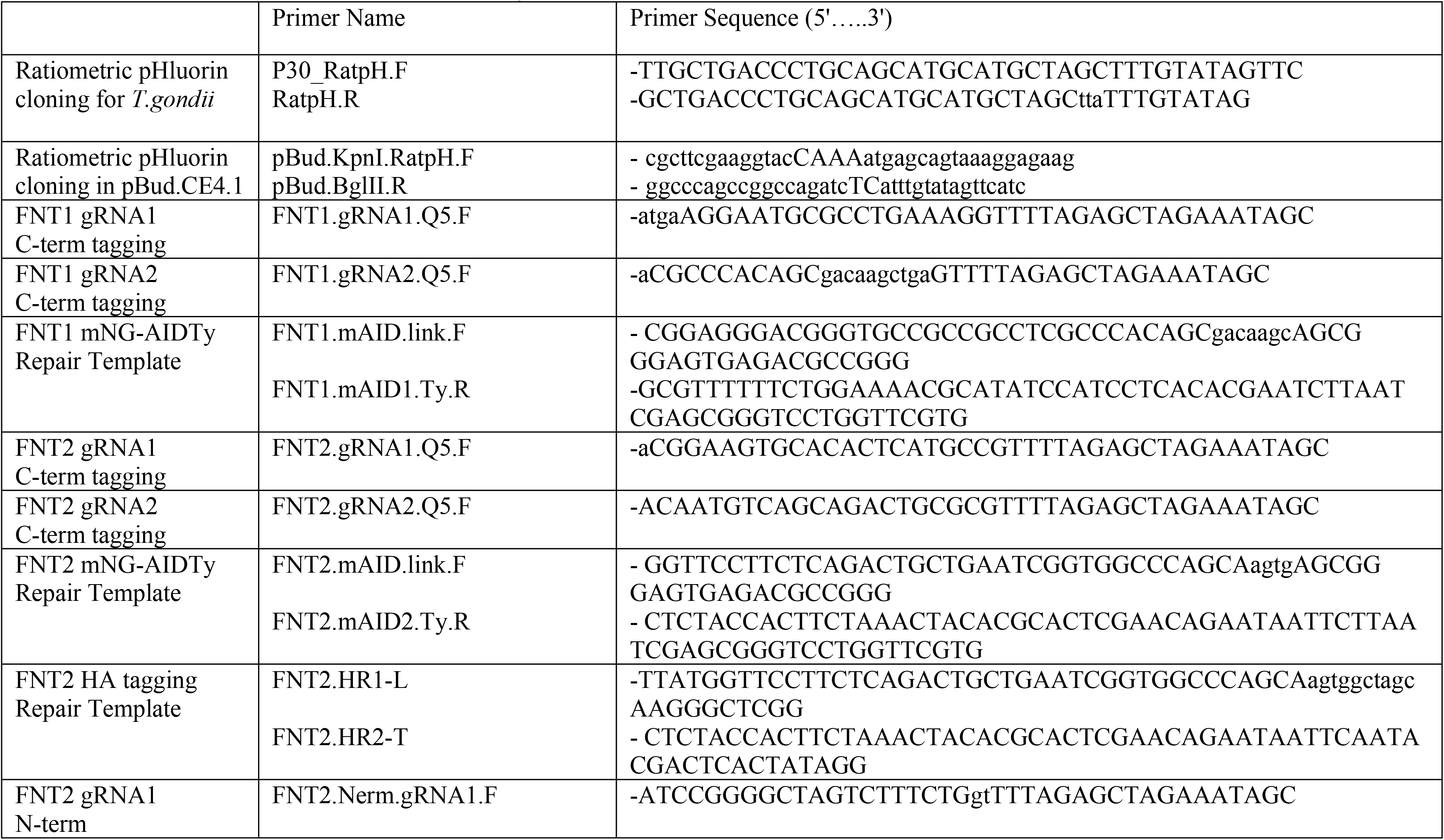
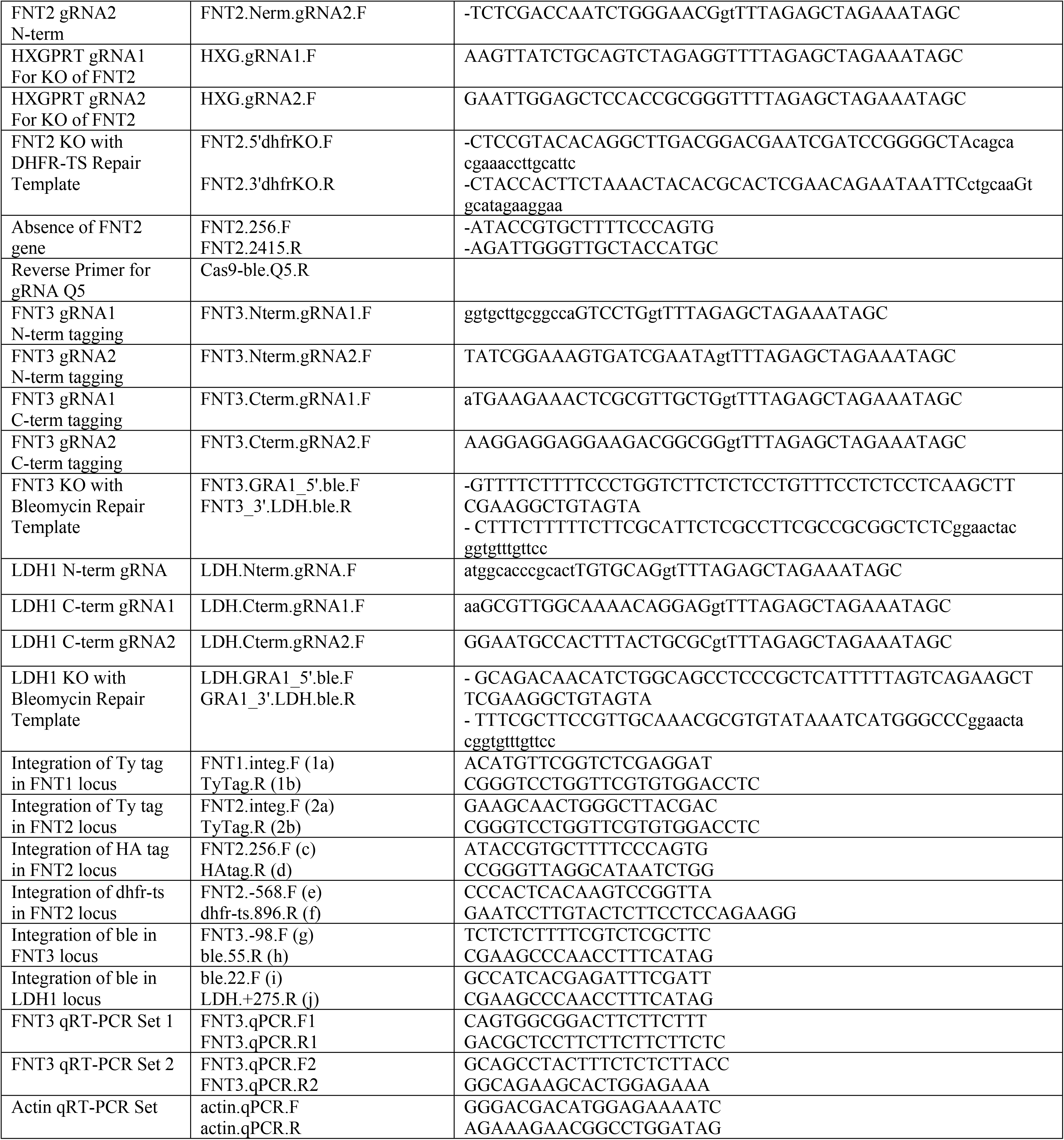
Primers used in this study.

The codon optimized pHluorin was amplified with primer P30_RatpH.F and Ratph.R and Gibson cloned into the *Avr*II restriction site of the P30_GFP_GPI plasmid (44), replacing the GFP. The RatpH plasmid was co-transfected with a DHFR-TS selectable plasmid (45) into RH parasites and selected with 1 μM pyrimethamine. Individual clones were isolated by limited dilution in 96-well plates, and one validated clone was selected and termed RH-RatpH. RH-RatpH was transfected with a red genetically encoded Ca^2+^ indicator (jRGECO1a in pCTH3, kindly provided by Dr. Silvia Moreno, University of Georgia) and an individual clone was isolated following selection with 20 μM chloramphenicol. These parasites are termed RH-RatpH:RGECO.

All mutant strains not stably expressing RatpH were transfected with 75-100 μg of RatpH and transiently transfected vacuoles were used in live imaging and pH experiments 38-40 h post-transfection.

Ratiometric pHluorin (Addgene, VV064: 1xCox8 - ratiometric pHluorin in fck) optimized for expression in mammalian cells was amplified and cloned in the pBud.CE4.1 plasmid in the *Kpn*I and *Bgl*II sites. For HeLa transfections, ∼80% confluent HeLa cells in an 8-well Ibidi slide was transfected with 500ng of pBud.pHluorin with 1.5μl of Fugene (Promega) and 50μl of Opti-MEM. Cells were infected with RH-RatpH ∼40hrs post-transfection and imaged ∼28-30hrs post-infection.

FNT1 and FNT2 CRISPR gRNA constructs were generated for C-terminal tagging using the NEB Q5 Site Directed Mutagenesis kit according to Brown et al. (46). Two FNT1 gRNAs (FNT1.gRNA1.Q5.F and FNT1.gRNA2.Q5.F) and two FNT2 gRNAs (FNT2.gRNA1.Q5.F and FNT2.gRNA2.Q5.F) with amplified with Cas9-ble.Q5.R were introduced into the Cas9-ble vector (47). Repair templates containing 40 bp of homology at the 5’ end and at the 3’ end to FNT1 or FNT2 and flanking the mNeonGreen.mAID.Ty (mNG-AIDTy) cassette were amplified (48). For transfection, 20 μg of each repair template along with 10 μg of each of the two gRNA constructs were transfected into RHΔ*hxgprt*Δ*ku80*:TIR parasites (24). Upon lysis 2 days post-transfection, extracellular parasites were selected with a final concentration of 50 μg/ml phleomycin in DMEM for 6 h at 37°C and 5% CO_2_, washed with DMEM, and placed back into a T25 containing HFF cells. Parasites were then cloned into 96-well plates and tested for the integration of the mNG-AIDTy cassette by PCR with primers FNT1.integ.F and Ty.Tag.R for FNT1 and FNT2.integ.F and Ty.Tag.R for FNT2

FNT2 was tagged at the C-terminal end in *iFNT1* parasites using the same CRISPR-Cas9 gRNA constructs as used for introducing the mNG-AIDTy cassette. The repair template was amplified from the pLinker.6xHA.HXGPRT.LoxP plasmid (49) and consisted of the 6xHA and HXPGRT selectable marker. Twenty micrograms of the repair template was transfected into *iFNT1* parasites, followed by selection with mycophenolic acid/xanthine. Integration of the 6xHA tag into the FNT2 locus was detected by PCR with primers FNT2.256.F and HAtag.R). Individual clones were isolated by limiting dilution, resulting in the strain *FNT2*-HA.

FNT2 knockout: Two gRNAs targeting the N-terminal end of *FNT2* (FNT2.Nerm.gRNA1.F and FNT2.Nerm.gRNA2.F) and two gRNAs against the 3’ end of the 6xHA.HXGPRT cassette (HXG.gRNA1.F and HXG.gRNA2.F) were generated in the Cas9-ble vector. A repair template to knock out *FNT2* with the DHFR-TS selectable marker was amplified with primers FNT2.5’dhfrKO.F and FNT2.3’dhfrKO.R and transfected with all four gRNAs (two for FNT2 and two for HXGPRT). Transfected parasites were selected with 1 μM pyrimethamine and individual clones were isolated and tested for the integration of the DHFR-TS selectable marker into the *FNT2* locus with primers FNT2.-568.F and dhfr-ts.896.R and for the absence of the FNT2 gene with FNT2.256.F and FNT2.2415.R. Individual clones were isolated by limiting dilution, resulting in the strain *iFNT1*Δ*fnt*2.

An LDH1 knockout strain was generated with an N-terminal gRNA (LDH.Nterm.gRNA.F) and two C-terminal gRNAs (LDH.Cterm.gRNA1.F and LDH.Cterm.gRNA.2.F) and a repair template using LDH.GRA1_5’ble.F and GRA1_3’LDH.ble.R to amplify the bleomycin selectable cassette. The repair template and gRNAs-Cas9-ble were transfected into *iFNT1*Δ*fnt*2 and selected as described above, resulted in the strain *iFNT1*Δ*fnt*2Δ*ldh1*.

The *iFNT1*Δ*fnt*2Δ*fnt3* strain was generated in a similar method with N-terminal gRNAs (FNT3.Nterm.gRNA1.F and FNT3.Nterm.gRNA2.F) and C-terminal gRNAs (FNT3.Cterm.gRNA1.F and FNT3.Cterm.gRNA2.F) and repair template amplification with FNT3.GRA1_5’.ble.F and FNT3_3’.LDH.ble.R.

### Live imaging and induced or natural egress

HFF monolayers were plated onto Ibidi 8-well chamber slides (#1.5 polymer coverslip) in phenol-free DMEM/10% cosmic calf/20 mM HEPES and grown overnight. Parasites expressing pHluorin were inoculated into the slides the next day and allowed to grow for 28-30 h prior to imaging. Wells were washed with Ringer’s buffer to remove any traces of background observed with DMEM and placed in 150 μl of Ringer’s. Slides were placed in a heating chamber on a Zeiss AxioObserver at 37°C and 5% CO_2_. Solutions to be used were also kept at 37°C in a heating block.

Following equilibration of the slide to the temperature and environment of the chamber, the time course experiment was started, and 5 frames were taken as the baseline measurement. This is followed by the addition of 150 μl of Ringer’s containing 2x inducer (400 μM zaprinast or 1 μM ionomycin) with tubing. For cytochalasin D or mycalolide B experiments, wells were incubated in 2 μM of cytochalasin D or 3 μM of mycalolide B in Ringer’s for 2 min prior to addition of inducer.

For natural egress, infected Ibidi slides were infected with RH-RatpH and allowed to develop at 37°C and 5% CO_2_. Videos were collected following ∼48-52 h of incubation and one frame was taken every 20sec for 20min. A new vacuole was then observed due to photobleaching of the vacuole.

The settings for taking images were as follows: Binning at 2×2, gain of 2, 1 frame every 1.5-5 seconds (or as indicated on graphs), 150 ms exposures each of the 410 nm and 470 nm channels. Custom pHluorin filter cube (Chroma) with excitation 395-415 nm and emission 500-550 nm for the 410 nm measurement and Filter Set 38HE (Zeiss) with excitation 450-490 nm and emission 500-550 nm for the 470 nm measurement were used. For parasites that also express RGECO, an additional image using a filter cube with excitation 540-580 nm and emission at 593-668 nm were taken. Images were converted from Zeiss czi files and exported as tiff files.

### pH calibration

RatpH expressing parasites in Ibidi chamber slides were incubated with Ringer’s buffer and 30 μM nigericin at pHs ranging from 5.5 to 8.5 and allowed to equilibrate for 3 min. At each pH, a 410 nm measurement and a 470 nm measurement were taken. These values were used to generate a ratio of 410/470 nm, which was plotted against the known pH to create a pH calibration curve in Prism. The resultant linear regression equation was then applied to the experimental ratios to extrapolate a pH value.

### pH measurements in vacuoles

Each timed experiment at each channel was imported into ImageJ (one video at 410 nm, one at 470 nm, and one at 560 nm), converted to 32-bit, and a StackReg Plugin was used to align images. An individual vacuole in the 410 nm channel was selected with the FreeHand Selector and marked as a region of interest with ROI Manager. The Measure Stack Plugin was used to give the measurement of the Raw Integrated Density. This ROI was then measured in the 470 nm and the 560 nm channels (if the strain tested expresses RGECO). A ratio was generated by dividing the 410 nm value by the 470 mm, and a pH value was extrapolated using the pH calibration curve and the associated linear regression equation. The pH values for each vacuole were graphed in Prism and a drop in pH was registered as a change of more than 0.05.

In addition to a pH calibration curve, each experiment included a timed experiment using DMSO in place of an egress inducer. This provides a measurement of the changes in pH from photobleaching of one of the channels. If it appeared that the pH trace changed/dropped through the time course, then the experimental traces were normalized by fitting a linear regression curve of the slope of DMSO trace.

### Buffer composition pH measurements

Chamber slides inoculated with RH_RatpH parasites were grown in phenol-free DMEM for 28-30 h as described above. Prior to imaging, chamber wells were rinsed with warm Ringer’s buffer and then incubated with the test buffer for 5 min. Induced egress and timed experiments were then started as described above. All buffers used are found in **Table 1**.

### FNT inhibitors

BH-296 and BH-388 were generously provided by Drs. Holger Erler and Eric Beitz (University of Kiel). Stock solutions of 10 mM were prepared in DMSO, frozen in 5 μl aliquots, and diluted to final working solutions of 1 μM, 5 μM, or 10 μM in Ringer’s buffer.

### Auxin-induced depletion of mAID-tagged proteins

A stock of 500 mM of indole-3-acetic acid (IAA) in 100% ethanol was diluted 1:1000 to knockdown expression of mAID-tagged proteins. Parasites were incubated in 500 μM IAA for 1 h, 2 h, and 6 h prior to isolation of the lysates for western blot and probing with RbαTy antibody. Similarly, intracellular parasites in chamber slides were incubated with 500 μM of IAA and observed live for the loss of mNG or fixed and IFAs were performed using RbαTy antibody.

### Immunofluorescence microscopy

HFF monolayers on glass chamber slides were infected with parasite strains and fixed with 4% paraformaldehyde. Slides were permeabilized with 0.1% Triton X-100, blocked with 5% FBS and 5% BSA, and probed with primary antibody in Wash Buffer (WB, PBS/1% FBS). Slides were then washed with WB and incubated with secondary antibody in WB. Following washes, slides were mounted with Mowiol and viewed on a Zeiss AxioObserver. RbαTy 1:1000, RbαGAP45 1:1000, MsαHA 1:1000, MsαIMC 1:500,

### Western blotting

Parasites were filter-purified, chased with cold PBS, pelleted at 1,000g at 4°C, washed with cold PBS, and centrifuged again. Parasite pellets were resuspended in RIPA buffer (50 mM Tris-HCl [pH 7.5], 1% NP-40, 0.5% sodium deoxycholate, and 0.1% SDS, 150 mM NaCl) and incubated at room temperature for 10 min. Lysates were centrifuged at 15,000g at 4°C for 10 min and the supernatant was removed. Room temperature 4x NuPAGE LDS sample buffer was added to the supernatant, along with 2-beta-mercaptoethanol to a final concentration of 2%. Lysates were separated on SDS-PAGE gels and semi-dry blotted onto PVDF or nitrocellulose membranes. Membranes were blocked with 5% milk in PBS-Tween, incubated with antibodies diluted in 1.25% milk in PBS-Tween, and visualized with West Pico ECL substrate (Thermo Scientific).

### Statistical Analyses

All statistical analyses were performed in GraphPad Prism. For each set of data, outliers were identified and removed using the ROUT method and an aggressive Q value of 0.1%. The data were then tested for normality and lognormality using the D’Agostino-Pearson test. If the data passed, a parametric One-way ANOVA with Tukey’s multiple comparisons test was used. If any set of data failed the normality test, a non-parametric Kruskal-Wallis test was used. Results were corroborated by a Mixed Effect Model generated from an RStudio script written by CSCAR (Consulting for Statistics, Computing, and Analytics Research) at the University of Michigan and considers random effects of the experiment. The macro/script is provided in Supplemental Information.

### L-Lactate and Pyruvate Detection Assay

Parasites with and without IAA treatment for 48 h was purified through a 3 μM filter, chased with Ringer’s buffer, and resuspended to 4×10^8^/ml. One hundred μl of each strain and treatment were placed into one well of a round-bottom 96-well plate and floated in a 37°C water bath. To each well, 100 μl of prewarmed 400 μM zaprinast (2X) was added and incubated for 10 min. The plate was then moved on ice for 5 min, and then centrifuged at 4°C for 10 min. After centrifugation, 175 μl of the supernatant (ESA) was removed from each well, placed in another well, re-centrifuged, and 150 μl was then removed. For each lactate detection assay well, 25 μl of the ESA was used. Lactate detection was performed following manufacturer’s protocol, Promega Lactate-Glo (Cat.# J5021), in a white 96-well plate and read on a BioTek Synergy H1 plate reader, Luminescence endpoint, Gain 135.

For the pyruvate assay, 20 μl of the ESA supernatant was tested following the manufacturer’s protocol, Cayman Chemical (Cat.# 700470). Black plates were read on a BioTek Synergy H1 plate reader, with excitation at 530 nm and emission at 590 nm.

### Reverse transcriptase qPCR

RNA extraction of parasites was performed using Qiagen RNEasy columns, followed by cDNA synthesis with Invitrogen Superscript III First-Strand Synthesis. cDNA products were used with SYBR Green Mix using the following reaction conditions: 5 min at 95°C, then 10 sec at 95°, 30 sec at 60°, 5 sec at 65°C x 39 cycles. The fold change in relative expression was calculated using actin as a housekeeping gene for normalization.

## Acknowledgements

We thank AJ Stasic, Stephen Vella, and Silvia Moreno for the *iΔvha1* parasites and RGECO plasmid, Sebastian Lourido for the mNG-AIDTy plasmid, Gustavo Arrizabalaga for the Δ*nhe1* and Δ*nhe3* parasites, and Holger Erler and Eric Beitz for the FNT inhibitors. We appreciate the Consulting for Statistics, Computing, and Analytics Research (CSCAR) center at the University of Michigan. We thank Aric Schultz, Marijo Roiko, and Alfredo Guerra for critically reading the manuscript and members of the Carruthers lab for helpful discussions. Aric Schultz also helped to generate the GIF in Supplemental Figure S2. Figure 7 was created with BioRender.

## Funding Information

This work was supported by operating grants from the United States National Institutes of Health Grant (R01AI046675 to V.B.C.). The funding agency did not play a role in study design, data collection or analysis, or the decision to submit the work for publication.

## Supporting Information

**Supplemental Figure S1.** A) 410nM image of HeLa cells transfected with RatpH. Arrowhead indicates a transfected HeLa cell and arrow indicates a transfected and infected (with RH-RatpH) HeLa cell. B) Ratio image (410/470nM) of A), dashed lines indicate the vacuoles. C) Time-course of ratio images following ionomycin induction; time post-induction is indicated in the upper left corner of each image. D, E) pH tracings of infected and uninfected-transfected host cells or vacuoles induced with DMSO or ionomycin (D) or zaprinast (E).

**Supplemental Figure S2.** Time-lapse imaging of intracellular Ca^2+^ levels (RGECO fluorescence; upper) and pH (pHluorin ratio, lower) following zaprinast induction. Graphs to the right indicate the relative fluorescence units for Ca^2+^ and the pH values for pHluorin, corresponding to the time-lapse images on the left.

**Supplemental Figure S3.** PCRs to confirm knockouts of NHE1 and NHE3. A) Primers designed to amplify a product only when a knockout of NHE1 has occurred (as used in (Arrizabalaga et al., 2004). B) Primers designed against the NHE3 gene detects NHE3 in WT but not Δ*nhe3* parasites.

**Supplemental Figure S4.** NHE4-AIDTy localization and protein down-regulation. A) TIR or NHE4-AIDTy parasites grown in chamber slides for 24 h were fixed and stained with anti-Ty and anti-GAP45 antibodies. B) NHE4-AIDTy parasites treated for 24 h with or without IAA were purified and immunoblotted with anti-Ty. Immunoblotting with anti-tubulin was used as a loading control.

**Supplemental Figure S5.** A) Schematic diagram of the strategies used to endogenously tag or knock out genes, and the lineage. B) PCRs to detect integration of epitope tags or selectable markers as indicated by the primers (shown as letters) in A.

**Supplemental Figure S6.** A) FNT1-AIDTy parasites were inoculated into Ibidi chamber slides and treated with or without IAA for the indicated times. The mNG fused to FNT1 was visualized with live imaging using a YFP filter cube. B) TIR and FNT1-AIDTy parasites were treated for the indicated times with or without IAA, filter purified, and lysates immunoblotted with anti-Ty. Arrow indicates FNT1-AIDTy.

**Supplemental Figure S7.** Reverse transcriptase qPCR to detect FNT3 mRNA expression. Two separate primer sets were used. Data represent 5 biological replicates each with triplicates samples. Error bars are mean ± S.E.M.

**Supplemental Figure S8.** Replication Assays. Parasites were inoculated into 8-well chamber slides and allowed to replicate for 24 h prior to enumeration. A minimum of 250 PVs were counted. Data represent 3 biological replicates each with triplicate samples. Error bars are mean ± S.E.M.

**Supplemental Figure S9.** Impairing Na^+^/H^+^ exchangers attenuates PV acidification. RH-RatpH vacuoles incubated with or without amiloride. Data points represent changes in PV pH starting from baseline to a drop greater than 0.05 following induction with either ionomycin or zaprinast.

## Notes

### Competing Interest Statement

The authors have declared no competing interest.

## References

1. Bisio H, Soldati-Favre D. Signaling Cascades Governing Entry into and Exit from Host Cells by Toxoplasma gondii. Annu Rev Microbiol. 2019 Sep 8;73:579–99.

2. Borges-Pereira L, Budu A, McKnight CA, Moore CA, Vella SA, Hortua Triana MA, et al. Calcium Signaling throughout the Toxoplasma gondii Lytic Cycle: A STUDY USING GENETICALLY ENCODED CALCIUM INDICATORS. J Biol Chem. 2015 Nov 6;290(45):26914–26.

3. Bisio H, Lunghi M, Brochet M, Soldati-Favre D. Phosphatidic acid governs natural egress in Toxoplasma gondii via a guanylate cyclase receptor platform. Nat Microbiol. 2019 Mar;4(3):420–8.

4. Jia Y, Marq J-B, Bisio H, Jacot D, Mueller C, Yu L, et al. Crosstalk between PKA and PKG controls pH-dependent host cell egress of Toxoplasma gondii. EMBO J. 2017 02;36(21):3250–67.

5. Uboldi AD, Wilde M-L, McRae EA, Stewart RJ, Dagley LF, Yang L, et al. Protein kinase A negatively regulates Ca2+ signalling in Toxoplasma gondii. PLoS Biol. 2018 Sep;16(9):e2005642.

6. Endo T, Tokuda H, Yagita K, Koyama T. Effects of extracellular potassium on acid release and motility initiation in Toxoplasma gondii. J Protozool. 1987;34(3):291–5.

7. Roiko MS, Svezhova N, Carruthers VB. Acidification Activates Toxoplasma gondii Motility and Egress by Enhancing Protein Secretion and Cytolytic Activity. PLoS Pathog. 2014 Nov 6;10(11):e1004488.

8. Beauregard KE, Lee K-D, Collier RJ, Swanson JA. pH-dependent Perforation of Macrophage Phagosomes by Listeriolysin O from Listeria monocytogenes. J Exp Med. 1997 Oct 6;186(7):1159–63.

9. Geoffroy C, Gaillard JL, Alouf JE, Berche P. Purification, characterization, and toxicity of the sulfhydryl-activated hemolysin listeriolysin O from Listeria monocytogenes. Infect Immun. 1987 Jul;55(7):1641–6.

10. Noronha FS, Cruz JS, Beirão PS, Horta MF. Macrophage damage by Leishmania amazonensis cytolysin: evidence of pore formation on cell membrane. Infect Immun. 2000 Aug;68(8):4578–84.

11. Andrews NW, Abrams CK, Slatin SL, Griffiths G. A T. cruzi-secreted protein immunologically related to the complement component C9: Evidence for membrane pore-forming activity at low pH. Cell. 1990 Jun;61(7):1277–87.

12. Manceva S. Effect of pH and ionic strength on the cytolytic toxin Cyt1A: a fluorescence spectroscopy study. Biochim Biophys Acta BBA - Proteins Proteomics. 2004 Jun;1699(1–2):123–30.

13. Risco-Castillo V, Topcu S, Marinach C, Manzoni G, Bigorgne AE, Briquet S, et al. Malaria Sporozoites Traverse Host Cells within Transient Vacuoles. Cell Host Microbe. 2015 Nov 11;18(5):593–603.

14. Miesenbock G, Angelis DAD, Rothman JE. Visualizing secretion and synaptic transmission with pH-sensitive green fluorescent proteins. Nature. 1998 Jul 9;394(6689):192–5.

15. Lourido S, Tang K, Sibley LD. Distinct signalling pathways control Toxoplasma egress and host-cell invasion. EMBO J. 2012 Dec 12;31(24):4524–34.

16. Moudy R, Manning TJ, Beckers CJ. The loss of cytoplasmic potassium upon host cell breakdown triggers egress of Toxoplasma gondii. J Biol Chem. 2001;276(44):41492–501.

17. Vella SA, Moore CA, Li Z-H, Hortua Triana MA, Potapenko E, Moreno SNJ. The role of potassium and host calcium signaling in Toxoplasma gondii egress. Cell Calcium. 2021 Mar;94:102337.

18. Kafsack BF, Pena JD, Coppens I, Ravindran S, Boothroyd JC, Carruthers VB. Rapid membrane disruption by a perforin-like protein facilitates parasite exit from host cells. Science. 2009 Jan 23;323(5913):530–3.

19. Stasic AJ, Chasen NM, Dykes EJ, Vella SA, Asady B, Starai VJ, et al. The Toxoplasma Vacuolar H+-ATPase Regulates Intracellular pH and Impacts the Maturation of Essential Secretory Proteins. Cell Rep. 2019 14;27(7):2132–2146.e7.

20. Arrizabalaga G, Ruiz F, Moreno S, Boothroyd JC. Ionophore-resistant mutant of Toxoplasma gondii reveals involvement of a sodium/hydrogen exchanger in calcium regulation. J Cell Biol. 2004 Jun 7;165(5):653–62.

21. Barylyuk K, Koreny L, Ke H, Butterworth S, Lassadi I, Mourier T, et al. Global mapping of protein subcellular location in apicomplexans: the parasite as we’ve never seen it before. Access Microbiol [Internet]. 2019 Mar 1 [cited 2020 Apr 17];1(1A). Available from: https://www.microbiologyresearch.org/content/journal/acmi/10.1099/acmi.ac2019.po0252

22. Karasov AO, Boothroyd JC, Arrizabalaga G. Identification and disruption of a rhoptry-localized homologue of sodium hydrogen exchangers in Toxoplasma gondii. Int J Parasitol. 2005 Mar;35(3):285–91.

23. Francia ME, Wicher S, Pace DA, Sullivan J, Moreno SNJ, Arrizabalaga G. A Toxoplasma gondii protein with homology to intracellular type Na+/H+ Exchangers is important for osmoregulation and invasion. Exp Cell Res. 2011 Jun 10;317(10):1382--1396.

24. Brown KM, Long S, Sibley LD. Plasma Membrane Association by N-Acylation Governs PKG Function in Toxoplasma gondii. mBio. 2017 May 2;8(3):10.1128/mBio.00375-17.

25. Marchetti RV, Lehane AM, Shafik SH, Winterberg M, Martin RE, Kirk K. A lactate and formate transporter in the intraerythrocytic malaria parasite, Plasmodium falciparum. Nat Commun. 2015 Mar 31;6:6721.

26. Wu B, Rambow J, Bock S, Holm-Bertelsen J, Wiechert M, Soares AB, et al. Identity of a Plasmodium lactate/H(+) symporter structurally unrelated to human transporters. Nat Commun. 2015 Feb 11;6:6284.

27. Erler H, Ren B, Gupta N, Beitz E. The intracellular parasite Toxoplasma gondii harbors three druggable FNT-type formate and l-lactate transporters in the plasma membrane. J Biol Chem. 2018 09;293(45):17622–30.

28. Kloehn J, Oppenheim RD, Siddiqui G, De Bock P-J, Kumar Dogga S, Coute Y, et al. Multi-omics analysis delineates the distinct functions of sub-cellular acetyl-CoA pools in Toxoplasma gondii. BMC Biol. 2020 Jun 16;18(1):67.

29. Zeng JM, Hapuarachchi SV, Shafik SH, Martin RE, Kirk K, van Dooren GG, et al. Identifying the major lactate transporter of Toxoplasma gondii tachyzoites. Sci Rep. 2021 Dec;11(1):6787.

30. Abdelbaset AE, Fox BA, Karram MH, Abd Ellah MR, Bzik DJ, Igarashi M. Lactate dehydrogenase in Toxoplasma gondii controls virulence, bradyzoite differentiation, and chronic infection. Hakimi MA, editor. PLOS ONE. 2017 Mar 21;12(3):e0173745.

31. Yang S, Parmley F. Toxoplasma gondii expresses two distinct lactate dehydrogenase homologous genes during its life cycle in intermediate hosts. Gene. 1997 Jan;184(1):1–12.

32. Pomel S, Luk FCY, Beckers CJM. Host Cell Egress and Invasion Induce Marked Relocations of Glycolytic Enzymes in Toxoplasma gondii Tachyzoites. Soldati-Favre D, editor. PLoS Pathog. 2008 Oct 24;4(10):e1000188.

33. Xia N, Yang J, Ye S, Zhang L, Zhou Y, Zhao J, et al. Functional analysis of Toxoplasma lactate dehydrogenases suggests critical roles of lactate fermentation for parasite growth in vivo. Cell Microbiol. 2018;20(1).

34. Xia N, Zhou T, Liang X, Ye S, Zhao P, Yang J, et al. A Lactate Fermentation Mutant of Toxoplasma Stimulates Protective Immunity Against Acute and Chronic Toxoplasmosis. Front Immunol. 2018 Aug 10;9:1814.

35. Lavine MD, Arrizabalaga G. Exit from host cells by the pathogenic parasite Toxoplasma gondii does not require motility. Eukaryot Cell. 2008 Jan;7(1):131–40.

36. Morawska M, Ulrich HD. An expanded tool kit for the auxin-inducible degron system in budding yeast. Yeast. 2013 Sep;30(9):341–51.

37. Natsume T, Kiyomitsu T, Saga Y, Kanemaki MT. Rapid Protein Depletion in Human Cells by Auxin-Inducible Degron Tagging with Short Homology Donors. Cell Rep. 2016 Apr;15(1):210–8.

38. Nishimura K, Fukagawa T. An efficient method to generate conditional knockout cell lines for essential genes by combination of auxin-inducible degron tag and CRISPR/Cas9. Chromosome Res. 2017 Oct;25(3–4):253–60.

39. Modesti L, Danese A, Angela Maria Vitto V, Ramaccini D, Aguiari G, Gafà R, et al. Mitochondrial Ca2+ Signaling in Health, Disease and Therapy. Cells. 2021 May 25;10(6):1317.

40. Miranda K, Pace DA, Cintron R, Rodrigues JC, Fang J, Smith A, et al. Characterization of a novel organelle in Toxoplasma gondii with similar composition and function to the plant vacuole. Mol Microbiol. 2010 Jun;76(6):1358–75.

41. Schmidt JDR, Walloch P, Höger B, Beitz E. Aquaporins with lactate/lactic acid permeability at physiological pH conditions. Biochimie. 2021 Sep;188:7–11.

42. Ives HE, Yee VJ, Warnock DG. Mixed type inhibition of the renal Na+/H+ antiporter by Li+ and amiloride. Evidence for a modifier site. J Biol Chem. 1983 Aug 25;258(16):9710– 6.

43. Paris S, Pouysségur J. Biochemical characterization of the amiloride-sensitive Na+/H+ antiport in Chinese hamster lung fibroblasts. J Biol Chem. 1983 Mar 25;258(6):3503–8.

44. Weiss LM, Kim K. Toxoplasma gondii : the model apicomplexan : perspectives and methods. 1st ed. London: Academic Press; 2007. 777 p.

45. Donald RGK, Roos DS. Stable molecular transformation of Toxoplasma gondii: A selectable DHFR-TS marker based on drug resistance mutations in malaria. Proc Natl Acad Sci USA. 1993;90:11703–7.

46. Brown KM, Long S, Sibley LD. Conditional Knockdown of Proteins Using Auxin-inducible Degron (AID) Fusions in Toxoplasma gondii. Bio-Protoc. 2018 Feb 20;8(4).

47. Di Cristina M, Carruthers VB. New and emerging uses of CRISPR/Cas9 to genetically manipulate apicomplexan parasites. Parasitology. 2018 Aug;145(9):1119–26.

48. Harding CR, Gow M, Kang JH, Shortt E, Manalis SR, Meissner M, et al. Alveolar proteins stabilize cortical microtubules in Toxoplasma gondii. Nat Commun. 2019 23;10(1):401.

49. Long S, Brown KM, Drewry LL, Anthony B, Phan IQH, Sibley LD. Calmodulin-like proteins localized to the conoid regulate motility and cell invasion by Toxoplasma gondii. PLoS Pathog. 2017 May 5;13(5):e1006379.

